# Biomimetic hydrogel platform reveals active force transduction from retinal pigment epithelium to photoreceptors

**DOI:** 10.1101/2025.01.09.632113

**Authors:** Sanna Korpela, Maija Kauppila, Viivi Karema-Jokinen, Lassi Sukki, Pasi Kallio, Heli Skottman, Soile Nymark, Teemu O. Ihalainen

## Abstract

In the eye, the retinal pigment epithelium (RPE) maintains the functionality and welfare of retinal photoreceptors and forms a tight, interlocked structure with photoreceptor outer segments (POSs). The RPE-retina interaction is difficult to recapitulate *in vitro*, limiting the studies addressing the retinal maintenance functions of the RPE. To overcome this challenge, we constructed a retina-mimicking structure using a soft polyacrylamide hydrogel coated with Matrigel. This structure was introduced to RPE cells’ apical side to model the RPE-retina interface *in vitro*. As a result, RPE cells attached to the hydrogels during culture, enabling further studies of cell adhesion and force transduction between the RPE-hydrogel with rheology and traction force microscopy. These methods were applied to a critical interactive process between the retina and the RPE: phagocytosis of the aged tips of POSs enabling their renewal. During phagocytosis, RPE cells imposed considerable traction forces to the POS particles. The force generation was actin-dependent, and the forces were significantly reduced by the disruption of RPE’s actin cytoskeleton. These results add another layer to the diverse interaction mechanisms between the RPE and the neural retina and pave the way for further studies of the RPE-retina interplay.

## Introduction

Retinal pigment epithelium (RPE) is an epithelial monolayer located at the back of the eye beneath the retina. Unlike a typical epithelium, RPE cells do not face an empty lumen on their apical side but are structurally and functionally in physical contact with retinal photoreceptor outer segments (POS) (Lakkaraju et al., 2020; Strauss, 2005). The apical membrane extensions of RPE, known as microvilli, protrude between POS, forming an interlocked structure. This close proximity of the retina enables the RPE to efficiently perform one of its primary roles, retinal maintenance. RPE cells play several crucial roles in supporting retinal health, including formation of the blood-retinal barrier, nutrient and waste transportation between retina and underlying choroid, maintenance of ion homeostasis, growth factor secretion, and regeneration of the visual pigments (Lakkaraju et al., 2020; Strauss, 2005; Wimmers et al., 2007). In addition, RPE cells are responsible for the essential daily renewal of POS in a process called phagocytosis (Lakkaraju et al., 2020; Strauss, 2005; Young & Bok, 1969). The close interaction between the RPE and photoreceptors is mediated by interphotoreceptor matrix (IPM), a specialized and compartmentalized extracellular matrix (ECM), that participates in retina-RPE adhesion (Ishikawa et al., 2015; Johnson & Hageman, 1991; Lazarus & Hageman, 1992). In addition, IPM partakes in retinoid, oxygen and nutrient transport, and contains enzymes and growth factors. Unlike typical ECM, IPM lacks the collagen meshwork, and is mainly composed of glycosaminoglycans, proteoglycans, and hyaluronan, which are large, hydrophilic molecules. (Ishikawa et al., 2015; Mieziewska, 1996) The IPM physically bridges the RPE and POS, connecting the tissue layers together. Furthermore, RPE-retina adhesion is mediated by α_V_β_5_ integrins on the apical surface of RPE, intraocular pressure, metabolism, and net fluid transport from the retina (Ghazi & Green, 2002; Nandrot et al., 2008). Even though the details of the interactions’ strength and dynamics are not known, enzymatic degradation of IPM (Yao et al., 1990) and inhibition of IPM synthesis (Lazarus & Hageman, 1992) have been linked to retinal detachment, and IPM’s role in photoreceptor degenerative diseases has been discussed (Hagstrom et al., 1999; Ishikawa et al., 2015; Mieziewska, 1996).

The RPE-retina adhesion needs to be dynamic to allow RPE cells to internalize and digest parts of the POS in the daily phagocytosis process (Lakkaraju et al., 2020; Strauss, 2005; Young & Bok, 1969). The POS continuously grow from their base, and older tips of outer segments, which have participated in the phototransduction process for longer, are engulfed and recycled by the RPE. This recycling prevents the accumulation of photo-oxidative compounds to photoreceptor cells and IPM, and is vital for retinal welfare. (Moran et al., 2022) The internalization of POS particles by the RPE is a highly dynamic process, involving actin cytoskeleton reorganization and formation of phagocytic cups that scission parts of the POS for ingestion (Spitznas & Hogan, 1970; Umapathy et al., 2023). This ‘nibbling’ of the POS requires the interplay of actin and myosin motors and thus it highly likely leads to tensile or compressive force generation locally at the RPE cell membrane (Kwon & Freeman, 2020; Strick et al., 2009; Umapathy et al., 2023). However, whether forces are needed in the POS phagocytosis process and if the forces are transduced from the RPE to the retina is not known. While live cell imaging methods have recently enabled studying the dynamics of phagocytosis by RPE in more detail (Jiang et al., 2015; Toops et al., 2014; Umapathy et al., 2023), characterization of the RPE generated forces in both RPE-retina adhesion and phagocytosis remains a challenge. Furthermore, current RPE *in vitro* cell culture systems (Lakkaraju et al., 2020; Vaajasaari et al., 2011; Viheriälä et al., 2021) are missing the RPE-retina interface, limiting studies addressing the biophysical characteristics of the interaction. Based on the biomechanical properties of the retina (Ferrara et al., 2021), we hypothesized that the force transduction and biophysics of the RPE-retina interface could be modelled utilizing a soft hydrogel as a retina-mimicking structure. Bringing this hydrogel into contact with the RPE apical side would allow the RPE cells to form connections and transmit forces to the hydrogel.

Hydrogels are versatile tools for mimicking extracellular environments in cell culture conditions (Caliari & Burdick, 2016). Here, we show that the biomimetic retina hydrogel platform can be constructed from elastic polyacrylamide (PA) hydrogel. PA gels are clear, synthetic, and non-degradable hydrogels consisting of cross-linked acrylamide polymer chain network, and their surface can be functionalized with extracellular matrix proteins to facilitate cell adhesion (Denisin & Pruitt, 2016; Tse & Engler, 2010; Wouters et al., 2016). We show that human embryonic stem cell (hESC) and human induced pluripotent stem cell (hiPSC) –derived RPE cells can be cultured with the biomimetic PA hydrogel on the apical side of RPE without changing the morphology or functionality of the cells. Also, as RPE cells adhered on the hydrogels on their apical side, we conducted further studies of the biophysical properties of the RPE-hydrogel interaction by using rheology and traction force microscopy. Our results indicate that the RPE-hydrogel adhesion is reversible and actin -mediated. Moreover, by embedding POS particles to the hydrogel, we were able to show that RPE cells exerted considerable forces to the retina-mimicking hydrogel in a scale of pN/µm^2^ during phagocytosis, which supports the emerging view of RPE being a mechanically active partner for the retina. In addition, our retina-mimicking structure offers an inexpensive, reproducible, and accessible tool for further biophysical modeling of the RPE-retina interface.

## Results

### A retina-mimicking hydrogel can be constructed from polyacrylamide

To study the RPE-retina interaction *in vitro*, we constructed a soft hydrogel to mechanically mimic the photoreceptors interacting with RPE (Figure 1A). Individual photoreceptor cells have been reported to have an elastic modulus (Young’s modulus, E) of 0.2 – 1.0 kPa (Zhang et al., 2023) and the whole photoreceptor layer approximately 26 kPa (Qu et al., 2018). Moreover, the IPM between RPE and the photoreceptors has a high prevalence of hydrophilic glycoproteins and proteoglycans (Ishikawa et al., 2015). Based on these properties we prepared a soft deformable PA hydrogel with E = 2.8 kPa which mimics the photoreceptor cell layer stiffness. PA hydrogels are resistant to protein binding and, therefore, non-adhesive also for cells without adhesive protein coating. Thus, we modeled the IPM layer by coating the PA hydrogel with Matrigel, a commercially available solubilized basal membrane, using L-DOPA treatment of the gel (Wouters et al., 2016). Kauppila et al. (2023) have shown that this L-DOPA and ECM coating on PA gel surface does not affect local stiffness values significantly. Thus, by combining the soft PA hydrogel and its coating with Matrigel proteins, we were able to construct a system which mimics the photoreceptor cell layer and has the important IPM components for cell adhesion. The retina-mimicking hydrogel was then used together with hESC- and hiPSC -derived RPE cells.

**Figure 1.**
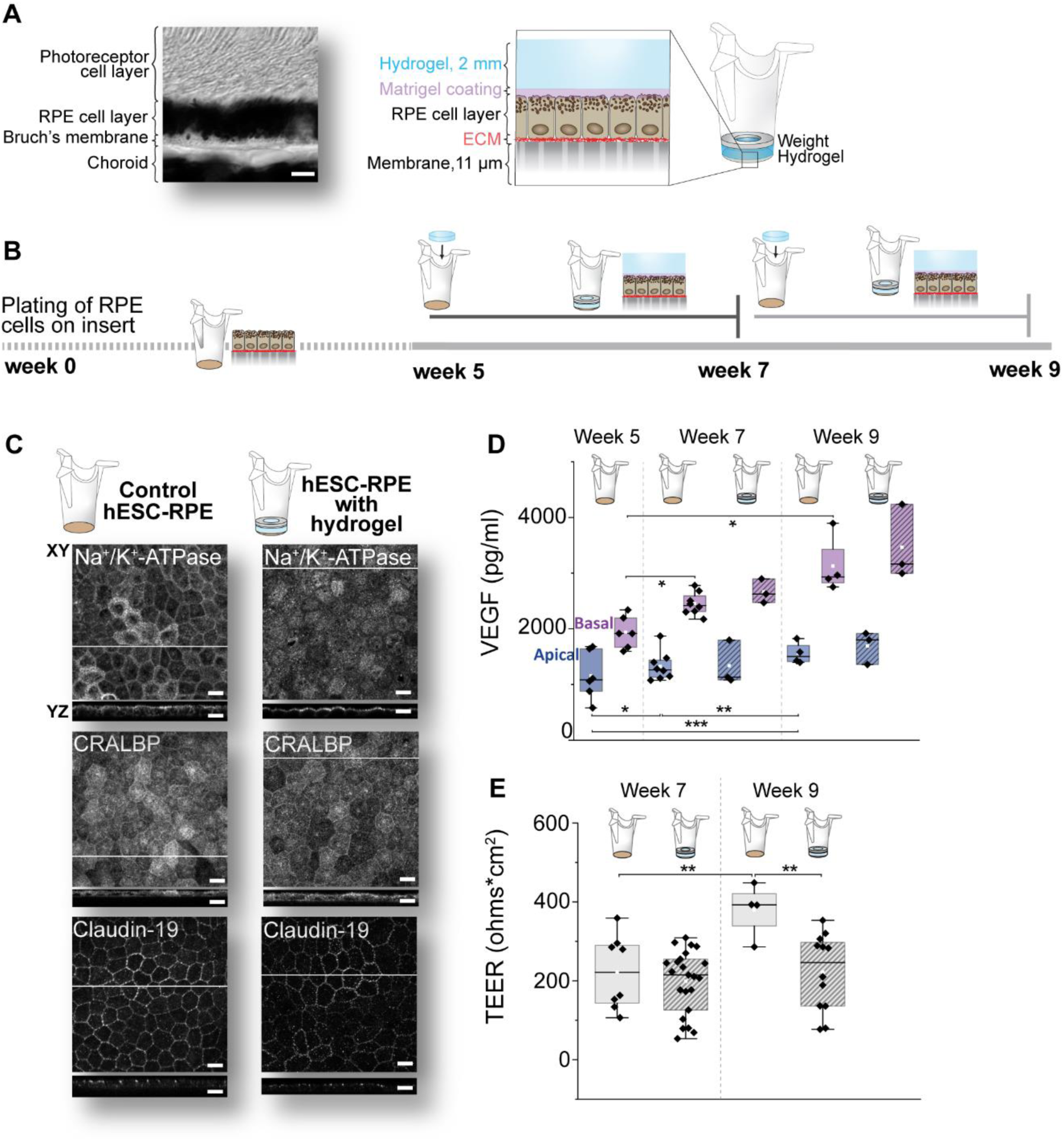
RPE can be cultured with retina-mimicking hydrogel without affecting its morphology or functionality. (A) Left: A brightfield image of a transverse section from mouse outer retina in paraffin, showing the close interaction between RPE cells’ apical side and photoreceptor cells. Scalebar 5 µm. Right: A schematic illustration of the RPE-hydrogel interaction. The PA hydrogel is coated with Matrigel and placed on the apical side of hESC- or hiPSC-derived RPE cells cultured in Transwell insert on a semipermeable membrane. During culture, RPE cells adhere to the hydrogel on their apical side. (B) Timeline of the cell culture experiments. hESC- and hiPSC-RPE cells were plated on 24-well inserts at week 0. After 5 weeks, the hydrogel was placed on the RPE apical side, and the cells were cultured with the hydrogel for two weeks until week 7. Alternatively, the hydrogels were placed on 7-week-old cells and cultured until RPE cells were 9 weeks old. (C) Confocal images of Na^+^/K^+^-ATPase, CRALBP, and claudin-19 RPE markers in 7-week-old hESC-RPE cells cultured without the hydrogel (control) and with the hydrogel for 2 weeks. hESC-RPE cells were labeled with immunofluorescence staining after removing the hydrogel. XY sections: maximum intensity projections. YZ sections: maximum intensity projections of 5 images with location indicated as a white line in the XY section. Scalebars 10 µm. (D) Vascular endothelial growth factor (VEGF) secretion was analyzed with ELISA assay from hESC-RPE cells cultured with the hydrogel for 2 weeks (controls cultured without the hydrogel). Data from two ELISA assays, with one datapoint in the plot containing multiple inserts (n = 41). Statistical testing with 2-way ANOVA with bootstrapping. (E) TEER was measured from hESC-RPE cells cultured 2 weeks with the hydrogel (controls cultured without the hydrogel). Statistical testing with 2-way ANOVA with Bonferroni correction. Significance codes: ‘*’ P < 0.05, ‘**’ P < 0.01 and ‘***’ P < 0.001.

First, the hESC- and hiPSC-RPE cells were plated on 24-well Transwell inserts (Figure 1B). After 5 weeks of culture, when RPE cells had formed a monolayer and started to form pigment but were still considered immature (Korkka et al., 2019; Vaajasaari et al., 2011), the hydrogels were placed on RPE apical side, allowing the cells to interact with the retina-mimicking hydrogel. The cells were cultured further with the hydrogel for 2 to 4 weeks. Control RPE cells were cultured without the hydrogel using the conventional culture method. Additionally, to test the hydrogel with more mature RPE, we used 7-week-old RPE cells and cultured them with the hydrogel until the cells were 9 weeks old. During culture, RPE cells were able to bind the Matrigel-coated PA hydrogels and adhered firmly to the hydrogel surface on their apical side.

Next, we wanted to investigate the hydrogels’ effect on RPE cell physiology and morphology. The hydrogels were carefully removed from the inserts, and cells were fixed with 4 % paraformaldehyde (PFA) and labelled with immunofluorescence staining. Labelling of the characteristic RPE markers Na^+^/K^+^-ATPase, CRALBP, and claudin-19 (Figure 1C) and Bestrophin, ZO-1, Ezrin, Ca_V_1.3, Connexin43, and vimentin (Figure 1 – figure supplement 1) showed no major differences between the RPE cells cultured under the hydrogel for 2 weeks and control RPE cells cultured without the hydrogel. To test one of the physiological functions of the RPE, vascular endothelial growth factor (VEGF) secretion was quantified with ELISA assay from both apical and basal side cell media (Figure 1D). VEGF secretion is known to be more active from the basal side of RPE *in vivo* (Korkka et al., 2019) and the same was observed in our RPE cultures with and without the hydrogel. In addition, VEGF secretion increased with time in control samples measured at week 5 (mean ± SD: apical side 1160 ± 430 and basal side 1940 ± 290 pg/ml), week 7 (apical 1330 ± 260 and basal 2450 ± 200 pg/ml), and week 9 (apical 1560 ± 200 and basal 3130 ± 520 pg/ml) after plating the cells (Figure 1D). However, RPE cells cultured with the hydrogel for 2 weeks showed no significant increase in the secreted VEGF between week 7 (apical 1340 ± 400 and basal 2660 ± 220 pg/ml) and week 9 (apical 1690 ± 290 and basal 3470 ± 670 pg/ml). Still, no significant differences at any time points were seen in VEGF secretion between controls and RPE-hydrogel samples. Finally, we followed the epithelial barrier formation under the gel by measuring transepithelial electrical resistance (TEER) values from control and RPE-hydrogel samples (Figure 1E). In control cells, TEER values increased significantly between weeks 7 and 9 (from 220 ± 90 to 380 ± 70 Ωcm^2^, mean ± SD). In RPE cells cultured with the hydrogel, TEER did not rise significantly (200 ± 80 Ωcm^2^ at week 7, and 220 ± 100 Ωcm^2^ at week 9). Thus, at week 9 control cells showed higher TEER values compared to RPE-hydrogel samples. However, both control and RPE-hydrogel samples showed cell-cell junction localization of claudin-19, an important RPE tight junction marker (Figure 1C) (Peng et al., 2011). This together with the measured TEER values indicate that both controls and RPE-hydrogel samples had functional tight junctions. All in all, RPE cells showed characteristic morphology as well as secretion and barrier functionality, demonstrating compatibility of the hydrogel with the RPE cell cultures.

The PA hydrogels used in this work have a flat surface, while *in vivo* RPE and POS form tight, interlocked connections with RPE microvilli protruding between the outer segments. Interestingly, we observed different RPE cell responses depending on whether the hydrogels were polymerized on glass coverslips or cell culture plastic. RPE cells cultured with hydrogels polymerized on glass showed abnormal and disrupted apical microvilli, whereas hydrogels polymerized on plastic led to apical actin structures in the RPE that were comparable to control cells without a hydrogel (Figure 1 – figure supplement 2). Glass coverslips have a smooth surface with a root mean square (RMS) roughness of less than 1 nm. In contrast, cell culture plastics have an RMS roughness of 1–6 nm, a factor known to influence cell adhesion and proliferation (Zeiger et al., 2013). We speculate that this nanoscale roughness is replicated on the hydrogel surface and contributes to the RPE response to the apically placed hydrogel. As a proof-of-concept, we also fabricated hydrogels with retina-mimicking topographical features on the PA hydrogel surface using polydimethylsiloxane (PDMS) molds created by photolithography. This approach successfully generated photoreceptor-sized wells on the hydrogel surface (Figure 1 – figure supplement 2, Appendix 1), potentially allowing RPE microvilli to protrude into these wells. However, the construction of these soft-topography hydrogels was technically challenging, and this work was continued using flat-surface PA hydrogels polymerized on cell culture plastics.

### RPE adheres heterogeneously to the hydrogel

Our results indicated that RPE can adhere to the apically placed hydrogel and that RPE cells’ physiology is not disturbed even during long culturing periods with the hydrogel. To further characterize the retina-mimicking system, the RPE-hydrogel samples were fixed with 4 % PFA and labeled with immunofluorescent staining, with the gel still adhered to the cells. Due to the transparency of the PA hydrogels, we were able to image the RPE cells through the hydrogel with a confocal microscope (Figure 2A). This allowed us to observe the RPE monolayer without disturbing the cell-hydrogel interface. The imaging of RPE cells without removing the hydrogel revealed typical tight junction structures labeled with ZO-1, apical membrane labeled with Ezrin, and actin cytoskeleton labeled with phalloidin. This indicates that the RPE can be easily observed through the hydrogel by using confocal microscopy.

**Figure 2.**
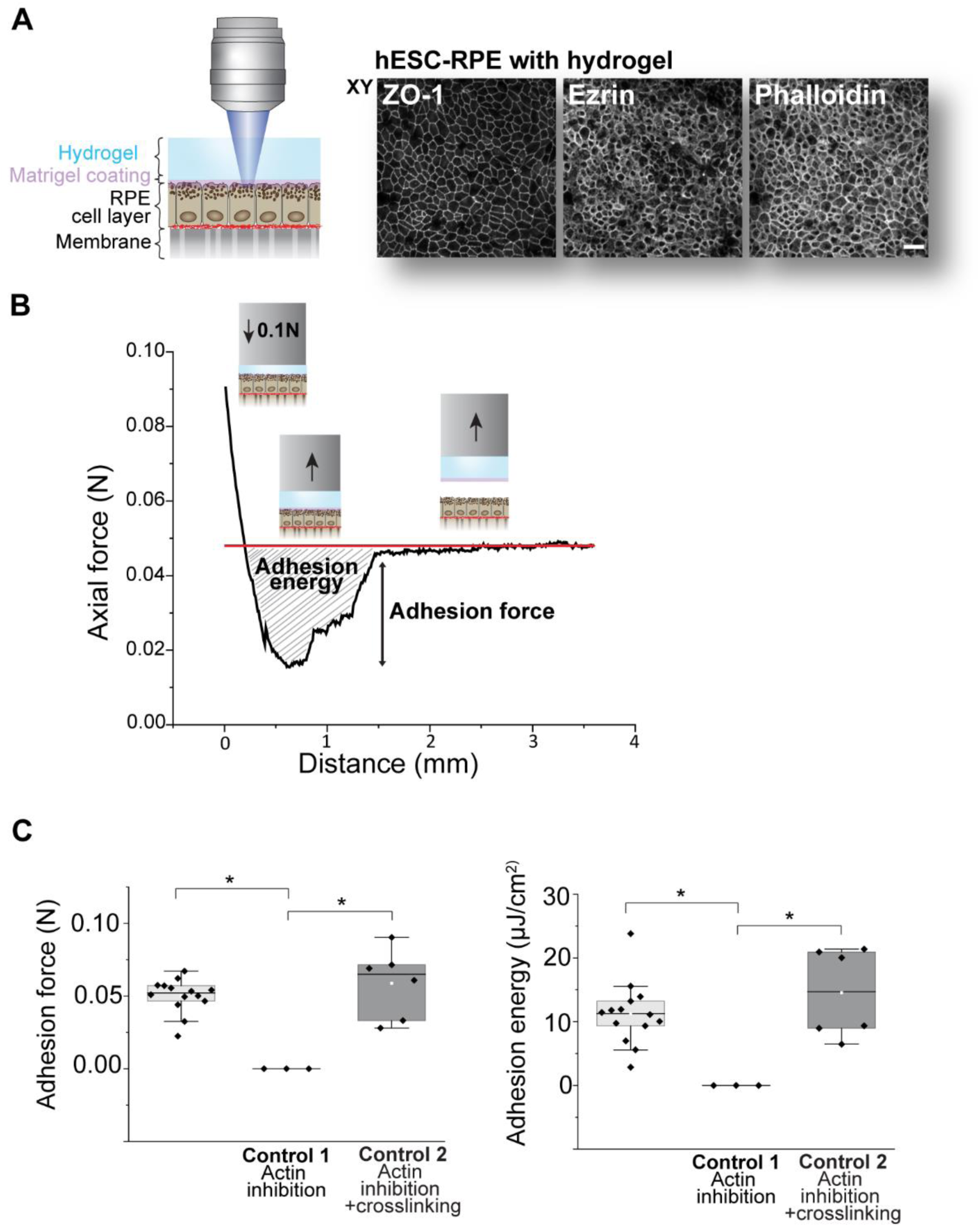
RPE-hydrogel adhesion was characterized with imaging and quantified with rheological tack adhesion test. (A) Schematic illustration of the imaging setup. RPE cells can be imaged through the hydrogel with a confocal microscope. hESC-RPE cells were fixed with 4% PFA and labeled with immunofluorescence staining with the hydrogel still attached to cells. ZO-1, Ezrin, and actin staining by Phalloidin. Scale bar 25 µm. (B) RPE-hydrogel adhesion was measured with a rheometric tack adhesion test. Live hESC-RPE cells with the hydrogel were placed on a rheometer and the probe was adhered to the hydrogel. Detachment was done by moving the probe up at a constant speed. The force-distance curve is a representative example of one measurement. At the end of the measurement, the force remained constant as the probe was moving upwards with the detached hydrogel still adhered to it. Adhesion force (maximum force measured, Newtons) was defined from each curve by subtracting this baseline force (indicated as a red line in the graph) from the lowest peak of the curve (detachment point). Adhesion energy (area over the curve, Joules per area) was measured graphically from each curve by calculating the area between the curve and the baseline force. (C) RPE-hydrogel adhesion force and energy were obtained from rheometric tack adhesion tests (n = 14 inserts). In control 1 (n = 3 inserts), RPE-hydrogel samples were treated with actin polymerization inhibitor Cytochalasin D for 1 hour, which detached the hydrogel from RPE cells, and the samples could not be measured (adhesion force and adhesion energy were = 0). For control 2 (n = 6 inserts), RPE-hydrogel samples were first treated with Cytochalasin D for 1 hour, and then fixed with 4 % PFA, which restored the adhesion. Statistical testing with Kruskal-Wallis. Significance codes: ‘*’ P < 0.05.

Previous experiments indicated that the RPE cells are able to adhere to the hydrogel and the hydrogel does not interfere the physiology of the RPE. To further characterize and quantify the adhesion between RPE and the hydrogel, a rheometric tack adhesion test was performed. The membrane with living RPE cells with the hydrogel was cut from the insert and glued on the bottom parallel plate of the rheometer. An 8 mm parallel plate was then adhered to the hydrogel with glue and maintained in 0.1 N compression for 2 min. Next, hydrogel detachment from the RPE cells was achieved by moving the probe upwards at a constant speed. As a result, a force-distance curve was obtained, where detachment of the hydrogel typically occurred stepwise instead of an instant detachment (Figure 2B). This indicates that the RPE-hydrogel adhesion is heterogeneous across the cell monolayer, with the weakest adhesion points detaching first. For each measurement, both maximum adhesion force and adhesion energy were determined from the force-distance curve (Figure 2C). After 2 weeks of culture, the average RPE-hydrogel adhesion force was 50 ± 10 mN (mean ± SD) per sample, corresponding to 1.7 ± 0.4 nN/µm^2^. Adhesion energy was 11.2 ± 4.9 µJ/cm^2^, corresponding to 112 fJ/µm^2^. As a first control, RPE-hydrogel samples were treated before the measurement with Cytochalasin D (10 µg/ml, 1 h), a pharmaceutical actin polymerization inhibitor. The treatment leads to depolymerization of actin fibers and a strong reduction of actomyosin contractility. Cytochalasin D caused complete spontaneous detachment of the hydrogels from RPE cells, corresponding to 0 adhesion force and energy. For a second control, RPE-hydrogel samples were first similarly incubated with Cytochalasin D but then treated with 4 % PFA, a cell fixative and protein crosslinker, which restored the adhesion to 50 ± 20 mN per sample, or 1.9 ± 0.7 nN/µm^2^. Adhesion energy was 14.5 ± 7.0 µJ/cm^2^, or 145 fJ/µm^2^. These controls suggest that the RPE-hydrogel adhesion is reversible and depends on intact actin cytoskeleton. However, actin depolymerization does not physically separate the hydrogel from the RPE in molecular scale, since they can be chemically crosslinked together after the depolymerization.

### RPE can transmit traction forces to the apical hydrogel

The previous experiments indicated that the RPE can adhere to the apically placed hydrogel and that the interaction does not influence the maturation of the RPE monolayer. The tight, actin-cytoskeleton-dependent adhesion thus mimics the close interaction between RPE cells and POS layer of the retina. Thus, the RPE-hydrogel adhesion enabled further studies of the RPE-photoreceptor mimicking interface. We hypothesized that the adhesion between these structures enable the direct transmission of RPE-generated forces from the RPE to the retina-mimicking hydrogel. We investigated this force transmission by conducting traction force microscopy (TFM). In TFM, fluorescent beads are embedded into the linearly elastic cell culturing substrate, e.g., hydrogel, and the forces transmitted across the adhesion between the cells and the substrate can be spatially quantified from the movement of the beads.

Here, we embedded fluorescent beads (diameter 200 nm) into the PA hydrogels prior to polymerization, placed the hydrogels on top of the RPE cells, and cultured for 2 or 4 weeks. RPE-hydrogel interface was then imaged twice with a 20 min interval through the hydrogel with a spinning disk confocal microscope to detect the bead displacements caused by the changes in cell-generated forces during the 20 min period (Figures 3A and 3B).

**Figure 3.**
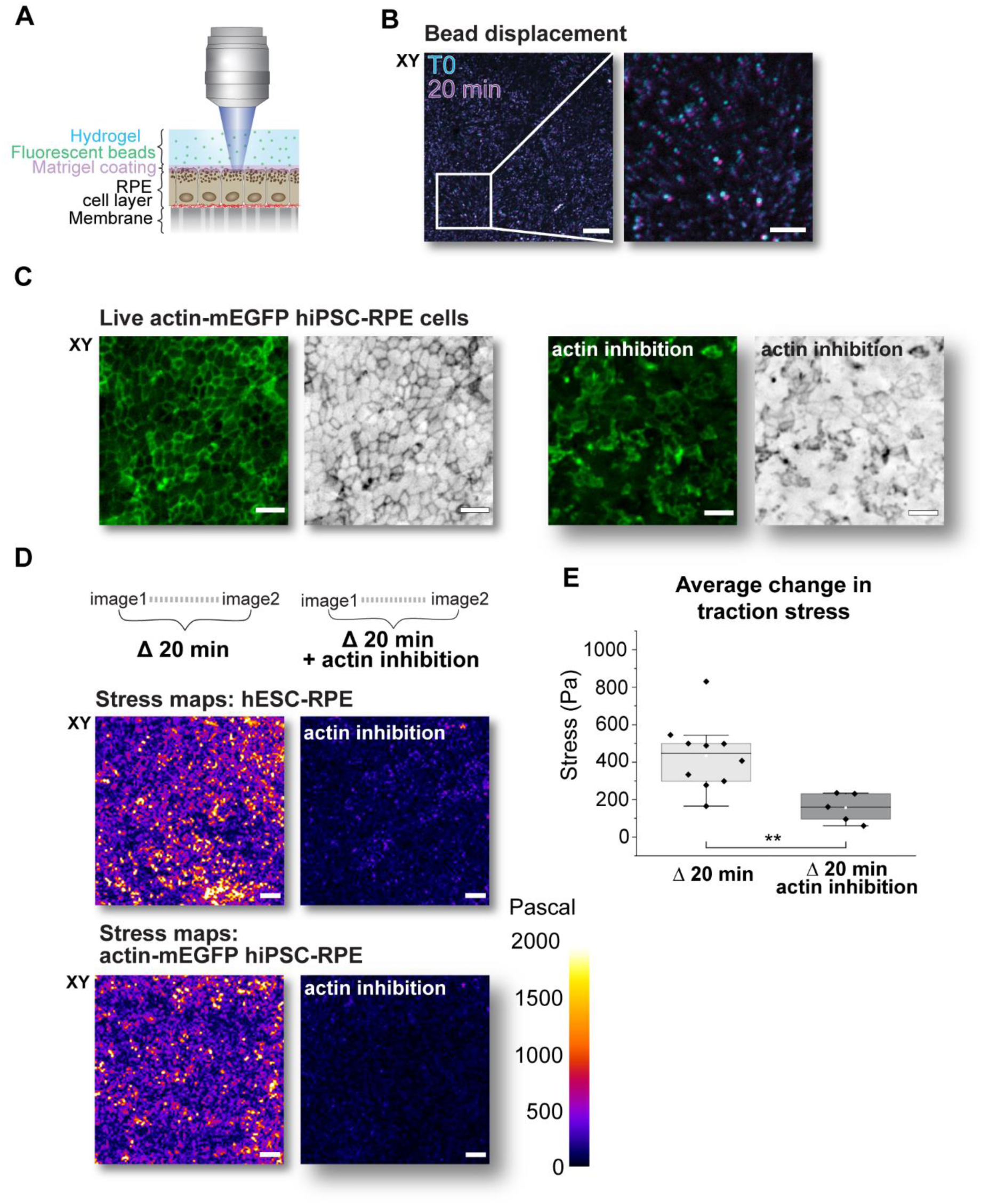
RPE-generated forces can be characterized with traction force microscopy. (A) Schematic illustration of the traction force microscopy (TFM) setup. The hydrogel is embedded with fluorescent beads, and live RPE-hydrogel interface is imaged through the hydrogel with spinning disk confocal microscope. Tractions exerted by RPE cells can be analyzed from the displacement of the beads. (B) Composite image of maximum intensity projections of the beads in RPE-hydrogel surface illustrating their displacement during a 20 min interval. Cyan: fluorescent beads at the start of the interval, magenta: beads after 20 min. Scalebars 50 µm (left) and 20 µm (right). (C) Live hiPSC-derived RPE cells with actin-mEGFP expression (green and inverted grayscale color) show that actin inhibition with Cytochalasin D affects actin cytoskeleton of the RPE cells with the hydrogel. Left: Untreated cells. Right: Cells after actin polymerization inhibition with 10 µg/ml of Cytochalasin D for 30 min. Scalebars 25 µm. (D) Bead displacements were calculated using particle image velocimetry (PIV) analysis and traction stress maps were constructed using Fourier transform traction cytometry (FTTC). Stress (Pascal) represents force per area (N/m^2^). Representative stress maps of hESC- and hiPSC -derived RPE cells’ interaction with the hydrogel, with and without Cytochalasin D. Scalebars 50 µm. (E) Average changes in cellular traction stresses over 20 min were calculated from each image pair. Data from 10 inserts, from which 5 inserts were used for actin inhibition with Cytochalasin D. Statistical testing with Mann-Whitney U. Significance codes: ‘*’ P < 0.05.

Our previous experiments indicated that the RPE-hydrogel adhesion can be significantly reduced with the actin polymerization inhibitor Cytochalasin D. To confirm that the Cytochalasin treatment leads to actin cytoskeleton disruption in the cells beneath the hydrogel, we generated hiPSC-RPE cells with endogenous beta-actin mEGFP expression (Figure 3 – figure supplement 1). This allowed us to follow actin dynamics in living cells under the hydrogel without additional labeling steps. Live imaging of actin-mEGFP hiPSC-RPE cells with the hydrogel showed disruptions in actin organization of the epithelial monolayer during the inhibition compared to the untreated cells (Figure 3C).

To compare the RPE-generated forces with and without an intact actin cytoskeleton, the beads inside the hydrogel were first imaged twice with a 20 min interval, and then RPE cells’ actin cytoskeleton was pharmaceutically depolymerized with Cytochalasin D for 40-60 min. The sample was then imaged for the second time, again twice with a 20 min interval. Bead displacement fields were computed from the microscopy image pairs taken 20 min apart with iterative particle image velocimetry (PIV) with a final vector spacing of 0.8 µm (Tseng et al., 2012). Since the rigidity and Poisson’s ratio of the PA hydrogels are known, the bead displacement fields were then used to construct traction vector plots (Figure 3 – figure supplement 2) and stress magnitude maps (Figure 3D) via Fourier transform traction cytometry (FTTC) (Tseng et al., 2012). The stress magnitude maps were heterogeneous and showed areas of low or high stress with local stress peaks, indicating variability in the activity of the RPE cells. The observed stresses generated by RPE tractions were diminished following the actin depolymerization with Cytochalasin D, indicating the importance of actomyosin generated forces in the RPE tractions.

For quantification, average values of the traction stresses were calculated from each image pair, corresponding to the average change in traction stress during the 20 min interval. With the hydrogel on their apical side, RPE cells generated lateral stresses on average of 430 ± 180 Pa (pN/µm^2^, mean ± SD) (Figure 3E). As can also be seen in the representative stress maps of both hiPSC- and hESC-RPE cells (Figure 3D), there was a decrease in traction stresses after inhibition of actin cytoskeleton with Cytochalasin D (on average to 160 ± 80 pN/µm^2^). This confirms that the bead displacements and thus the traction stresses analyzed here are caused by the actin cytoskeleton contractility of the RPE cells. These actin dependent forces are then transmitted to the retina-mimicking hydrogel on the apical side of the RPE cells via the cell-hydrogel contacts.

### RPE actively pulls on photoreceptor outer segments

RPE has an important role in the maintenance of the retina, and it actively recycles the tips of POS via phagocytosis. The process is strongly dependent on an intact actin cytoskeleton and integrin receptors on the apical surface of the RPE cells. (Lakkaraju et al., 2020) Since the RPE can transmit forces to the retina-mimicking hydrogel, we hypothesized that active force transduction happens during the phagocytosis process. This allows the RPE cells to pull the POS to the vicinity of the cell membrane, facilitating the internalization of the particles.

To study the forces that RPE cells generate during phagocytosis, thin PA hydrogels with fluorescent beads were constructed on glass coverslips. In addition, POS particles were extracted from porcine eyes, labeled with a fluorescent dye, and incorporated on the PA hydrogel surface. Covalent linkage and proper attachment of the POS particles to the PA gel was achieved by mixing NHS-acrylic acid to the PA gel precursor solution. This process yielded a soft hydrogel suitable for TFM, with POS particles on its surface. The hydrogel was not coated with any other ECM proteins, so the POS particles provided the only binding sites for the RPE cells. The insert membrane was then cut and placed on top of the hydrogel so that RPE cells faced down to the POS particles (Figure 4A). The thin hydrogel allowed imaging through both the glass coverslip and the hydrogel with a laser scanning confocal microscope to achieve better spatial resolution. We used SiR-actin to label the actin cytoskeleton of live hESC-RPE cells, allowing simultaneous imaging of the actin dynamics, bead displacement, and POS particles. During imaging, RPE cells were in close contact with the hydrogel, and RPE cells reached towards the POS particles with apical actin-rich - protrusions. Moreover, RPE cells were able to internalize parts of the POS particles incorporated in the hydrogel (Figures 4A and 4B).

**Figure 4.**
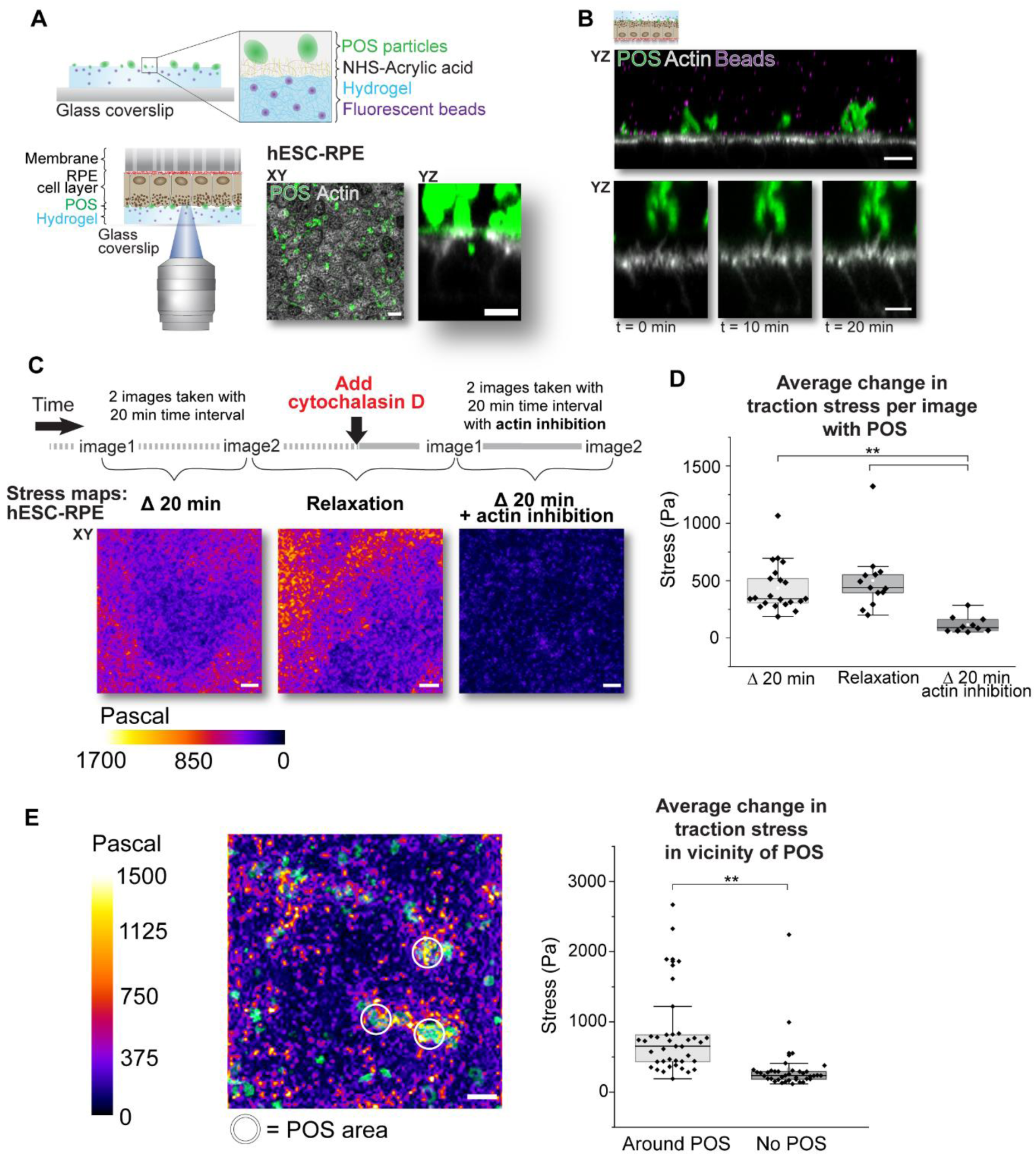
RPE actively generates forces during phagocytosis. (A) Schematic illustration of the imaging setup to analyze the force transduction during phagocytosis. Thin PA hydrogel (height approximately 100 µm) constructed on a glass coverslip was embedded with fluorescent beads and coated with NHS-acrylic acid and fluorescently labeled photoreceptor outer segment (POS) particles. Insert membrane with RPE cells was cut and placed on the hydrogel with RPE cells’ apical side facing towards the POS. Imaging was done through the glass coverslip and the thin hydrogel with a laser scanning confocal microscope. During imaging the RPE cells were kept in media supplemented with 10 % FBS. XY-maximum intensity projection shows RPE cells (grey, actin labeled with live cell dye SiR-actin) and POS particle localization above (green). Scale bar 10 µm. YZ image shows RPE cells from the side, with internalized POS particle inside the cell. Scale bar 5 µm. (B) Upper image: YZ-maximum intensity projection of 5 images from the RPE-hydrogel interface showing cellular actin labeled with SiR-actin (grey), fluorescently labeled POS particles (green), and beads embedded in the hydrogel (magenta). Scalebar 10 µm. Lower image: YZ-magnification of a single cell. Apical side microvilli (grey) protrude towards the POS particle (green) during 20 min. Scalebar 5 µm. (C) From each insert, 2-3 separate positions were chosen and imaged alternately. Traction stresses were analyzed from two images taken with an interval of 20 min (Δ20 min). 10 µg/ml of Cytochalasin D was added to the imaging media without moving the sample, and image acquisition in the positions was continued with 20 min intervals (Δ20 min with actin inhibition). In addition, analysis was performed by comparing the bead positions before and after the addition of Cytochalasin D (Relaxation). Average intensity Z-projections of FTTC stress maps from each condition (Δ20 min 21 images from 8 inserts, Relaxation 13 images from 5 inserts, and Δ20 min with actin inhibition 10 images from 5 inserts). Scalebars 10 µm. (D) From each image pair, an average value of the change in traction stresses during 20 min was calculated. Statistical testing with Kruskal-Wallis. (E) Left: Composite image of a traction stress map during cell cytoskeleton relaxation with maximum intensity Z-projection of POS particles (green). The white circles illustrate the areas (d = 10 µm) around the POS particles chosen for the analysis. Scale bar 10 µm. Right: Average traction stresses were calculated from each chosen area in vicinity of POS particles (n = 39 positions from 13 images, from 5 inserts). Control areas were randomized areas within the same images. Statistical testing with Mann-Whitney U. Significance codes: ‘**’ P < 0.01 and ‘ns’ P > 0.05.

Images were again acquired at 20 min intervals from different positions of the sample (Figure 4C). Next, Cytochalasin D (10 µg/ml) was added to the imaging media without moving the sample and incubated for at least 60 min. The same positions of the sample were imaged again, and traction stresses were analyzed from the acquired data before and after actin inhibition. Finally, we compared two images taken before and after the addition of Cytochalasin D, to study the change in stress magnitude during complete cell cytoskeleton relaxation. Analysis was performed by first computing bead displacements with PIV analysis from image pairs with 20 min intervals, and then calculating the traction vector plots (Figure 4 – figure supplement 1) and stress maps with FTTC. Averages of the stress magnitude maps indicate that actin polymerization inhibition decreases traction stress generation in RPE during POS phagocytosis (Figure 4C).

For quantification, average values of the traction stresses were calculated from each image pair, corresponding to the average change in traction stress during the 20 min interval (Figure 4D). During phagocytosis, RPE cells generated stresses on average of 430 ± 200 Pa (pN/µm^2^, mean ± SD). Interestingly, tractions measured within 20 min intervals did not differ significantly from the ones during relaxation (500 ± 280 pN/µm^2^), suggesting a very dynamic force generation in RPE especially at the early phases of the phagocytosis process. The actin inhibition with Cytochalasin D decreased traction stress generation during the 20 min interval to 120 ± 70 pN/µm^2^. As a control, we used hydrogels without any surface modifications (Figure 4 – figure supplement 2). In the absence of POS particles, hydrogel’s NHS-acrylic acid supplementation, and serum the quantified traction forces were only a few tens of pN/µm². Furthermore, the stresses were again negligible with the addition of serum (10 % FBS) and NHS-acrylic acid, demonstrating that POS particles are essential for force transduction and subsequent traction stress, and that RPE cells are unable to attach to the gel without them.

Finally, we wanted to compare the magnitude of the traction stresses around POS particles to areas without POS (Figure 4E). Three circular areas around POS particles (approximately 10 µm in diameter) were chosen in each traction stress map, representing the change in traction stress during cell cytoskeleton relaxation. Control areas were identical areas randomly within the same image. Traction stresses were significantly higher near the POS particles (800 ± 590 pN/µm^2^) compared to the randomly chosen control areas (320 ± 350 pN/µm^2^). This result indicates that the tractions measured here are mainly caused by RPE interacting with the POS particles on the hydrogel surface. Altogether, this data shows that RPE cells are able to focus physical force to the vicinity of POS during early phases of phagocytosis.

## Discussion

Close interaction between the retina and its underlying tissue, the retinal pigment epithelium (RPE), is vital for retinal wellbeing and is accompanied with tight adhesion between the RPE and the photoreceptor outer segments (POSs). As part of this, microvilli on the RPE apical side protrude between the POSs and form an interlocked, physical connection. This close connection is especially evident during daily phagocytosis of POS tips by the RPE (Lakkaraju et al., 2020; Strauss, 2005). We studied the physical characteristics of the photoreceptor-RPE interactions by developing a new hydrogel-based model system. Our results reveal that forces are transmitted from the RPE to the hydrogel and that the RPE cells exert actin cytoskeleton dependent physical forces to POS particles during their phagocytosis.

The present work shows that a retina-mimicking structure can be constructed from soft polyacrylamide (PA) hydrogel. Culturing of RPE with the apically residing gel did not lead to significant differences in cell morphology, RPE marker expression, epithelial layer integrity, or growth factor secretion, compared to cells cultured without a hydrogel (Figure 1). The measured TEER values of the RPE were around 200 Ωcm^2^, which is in line with the measurements from human tissue (Quinn & Miller, 1992) and porcine primary RPE cell cultures (Toops et al., 2014). Furthermore, the RPE cells adhered to the hydrogel from their apical side, enabling, to our knowledge for the first time, *in vitro* modeling of the physical RPE-retina interface.

Our model system allowed adhesion measurements between the RPE and the retina-mimicking hydrogel. The maximal RPE-hydrogel adhesion was approximately 170 mN/cm^2^, or 50 mN for an insert with 75000 cells (Figure 2). This corresponds to a gravitational force generated by a weight of 5.1 g or a vertical pulling force of approximately 17 bumblebees (Mountcastle & Combes, 2013). As a cell-level comparison, individual cells in endothelial monolayers have been reported to have a cell-substrate adhesion force of 1.17 µN measured with fluidic force microscopy, corresponding to approximately 90 mN for 75000 cells (Sancho et al., 2017). For physiologically relevant RPE-retina adhesion measurements *in vivo*, Chiang et al. (1995) showed that injection of Cytochalasin D, the same actin polymerization inhibitor we used in our experiments, into the subretinal space weakened the RPE-retina adhesion, and this effect was reversible. Thus, similarly to our *in vitro* model, intact actin cytoskeleton is needed for proper retina-RPE adhesion. However, when we fixed the Cytochalasin D treated samples with a paraformaldehyde crosslinker, we saw the restoration of the adhesion. This suggests that the adhesion is mediated by the RPE cells’ apical side and actin cytoskeleton depolymerization does not cause separation of the RPE and the hydrogel in the molecular-scale.

We investigated the force transduction with high spatial resolution using traction force microscopy (TFM). In TFM, fluorescent beads incorporated in the hydrogel are used to reconstruct the material deformation and subsequently the cell-generated traction forces leading to the deformations (Denisin et al., 2024). Our TFM experiments showed that after the hydrogel was cultured on the RPE apical side for 2-4 weeks, the RPE cells actively transmitted forces into the hydrogel (Figure 3). TFM images revealed a speckled pattern of force transmission, with some regions in the field of view showing higher tractions, indicating either heterogeneous adhesion between the RPE and the hydrogel or variability in RPE force production. Moreover, Cytochalasin D considerably decreased the tractions between RPE and the hydrogel, indicating that the forces are indeed rising from the actin cytoskeleton of the RPE cells. While the roles of integrin α_V_β_5_ (Nandrot et al., 2006) and actin (Chiang et al., 1995) in the RPE-retina adhesion have been addressed previously, our retina-mimicking hydrogel uniquely enabled quantification of mechanical force transmission from the apical side of RPE to the retina-mimicking hydrogel.

We next studied the possible force transmission from RPE to POS particles in phagocytosis. To quantify the traction forces that RPE cells generate during phagocytosis, we further developed the retina-mimicking hydrogel system. We constructed thin hydrogels on glass coverslips to enable imaging of the hydrogel surface with high-resolution confocal microscopy. The hydrogel was embedded with fluorescent beads to follow the RPE cell generated tractions, and purified POS particles from porcine eyes were incorporated on the hydrogel surface for RPE cells to phagocytose. During the experiments, RPE cells were able to contact POS particles and internalize parts of them. Previous studies have shown that RPE’s apical side actin-rich protrusions surround, incise, and ingest parts of the POS (Almedawar et al., 2020; Lieffrig et al., 2023; Matsumoto et al., 1987; Spitznas & Hogan, 1970; Umapathy et al., 2023) presumably with the help of myosin II (Strick et al., 2009; Zihni, 2025), suggesting that RPE cells can generate considerable forces during this process. We measured RPE generated traction forces in the scale of hundreds of pN/µm^2^ in the vicinity of the POS particles (Figure 4). As a comparison, the traction forces in individual focal adhesions in highly contractile cells can reach a few nN/µm², in line with our results during phagocytosis. Vorselen et al. (2021) used microparticle TFM to measure the force needed to deform 9 µm sized particles by macrophages during phagocytosis. The forces quantified from microparticle deformations increased from 1 nN to 10 nN in phagocytosis. The force magnitudes are reasonable considering the differences in the nature of the measured forces (pulling vs. deformation force) and cell types. In addition, our traction stress quantification by PIV relies solely on the in-plane movement of fluorescent beads, meaning that bead displacement directly toward or away from the RPE is not detected. Thus, the forces quantified here represent a lower estimation of the total magnitude of forces generated by the RPE, rather than measurement of total forces.

POS phagocytosis by the RPE is dependent on α_V_β_5_ integrin (Finnemann et al., 1997), and interestingly, α_V_β_5_-rich reticular adhesions have been shown not to associate with actin fibers (Lock et al., 2018). However, decrease in force generation after addition of Cytochalasin D shows that actin cytoskeleton participates in the RPE apical side force generation in phagocytosis. This is in line with previous work by Umapathy et al. (2023), where they showed with live cell imaging that during phagocytosis Cytochalasin D did not affect POS binding to the RPE apical surface but prevented the formation of actin-rich phagocytic cups.

Our results support the emerging view of RPE being a mechanically active participant in POS phagocytosis, rather than passively clearing shed POS particles from the subretinal space (Lakkaraju et al., 2020). This was achieved by developing a unique retina-mimicking hydrogel assay, which allowed us to study the mechanobiology of the RPE-retina interface. Currently there are no *in vitro* models which would recapitulate the interface of these two tissues. Organoids and organ-on-a-chip technologies offer interesting possibilities for *in vitro* culturing, but curiously, they are still missing the direct RPE-photoreceptor connections. While Gabriel et al. (2021) succeeded to generate RPE cells in optic vesicle-containing brain organoids, contacts between RPE cells and photoreceptors have generally been absent in retinal organoids (Chakrabarty et al., 2024; Zhao & Yan, 2024). Achberger et al. (2019) combined both RPE cells and retinal organoids in a microfluidic system to create a retina-on-a-chip. However, RPE and photoreceptors were still separated with hyaluronic acid -based hydrogel, thus restricting their direct connections. We chose PA for the retina-mimicking material based on its mechanical properties, well established polymerization and conjugation chemistry, and compatibility with high-resolution confocal microscopy. PA hydrogels are easy to construct at a low cost, and their elastic modulus can be tuned by simply changing the ratios of the monomer and the crosslinker, making them suitable for cell culture experiments (Denisin & Pruitt, 2016; Tse & Engler, 2010). In addition, cells cannot spontaneously adhere to PA hydrogels, but the surface can be modified by covalently linking selected ECM proteins using, for example, L-DOPA (Wouters et al., 2016). For the protein coating we used Matrigel, a commercially-available basement membrane ECM secreted by mouse sarcoma, that contains some of the main components of the IPM, such as laminin and heparan sulfate proteoglycans together with TGF-ß growth factors (Ishikawa et al., 2015).

Even though PA hydrogels are a simple and robust platform for cell culture experiments, more *in vivo* -like hydrogel materials could be considered for RPE cells, such as hyaluronic acid, one of the main components of the IPM, the specialized ECM found between RPE and photoreceptor cells in the eye (Ishikawa et al., 2015). However, mechanical properties, stability and the construction of the hyaluronic acid hydrogels would need to be tailored for the RPE cultures (Luo et al., 2023). Furthermore, for more *in vivo* -like retina-mimicking hydrogels, the surface topography of the gels could be considered. We showed that the RPE cell response differs between hydrogels polymerized on glass coverslips, compared to gels polymerized on cell culture plastics (Figure 1 – figure supplement 2). We speculate that the different nanoscale roughness of glass and plastic are transferred to the hydrogel surface during polymerization, leading to different RPE cell responses (Zeiger et al., 2013). Therefore, we constructed photoreceptor-sized wells to the PA hydrogel surface using replica molding (Appendix 1). However, construction of these wells on the soft PA hydrogel surface was challenging, and the outcome did not meet the requirements for our cell culture experiments. Therefore, future work should consider alternative materials and topographical fabrication techniques for more *in vivo* -like retina-mimicking platforms.

In conclusion, we have developed a retina-mimicking platform from PA hydrogel to be used with cultures of hESC- and hiPSC-derived RPE cells. The hydrogel platform enabled the direct measurements of the RPE adhesion to the retina-mimicking hydrogel without affecting RPE cell morphology or functionality. We showed, to our knowledge for the first time, that RPE cells exert considerable forces to POS particles during phagocytosis and that the force generation is dependent on an intact actin cytoskeleton. Together, our results support the emerging view of the RPE as a mechanically active partner for the retina and provide tools for further biophysical *in vitro* modeling of the RPE-retina interface.

## Materials and methods

### Polyacrylamide hydrogel construction

Polyacrylamide (PA) hydrogel construction was done based on the work of Tse & Engler (2010). For E = 2.8 kPa PA hydrogel, acrylamide (AA) (stock 40 %, #1610140, Bio-Rad Laboratories, USA) and Bis-acrylamide (Bis) (stock 2 %, #1610142, Bio-Rad Laboratories) were mixed in the ratio of AA 10 % and Bis 0.03 % in PBS in a falcon tube. Solution was degassed in vacuum. To initiate polymerization, 0.2 % v/v of TEMED (#1610800, Bio-Rad Laboratories) and 1 % v/v of ammonium persulfate (APS) stock solution in ddH_2_0 (10 % w/v, #A3678, Sigma-Aldrich, USA) were added to the PA solution and mixed by tilting the tube couple of times. The PA hydrogel polymerization was done by pipetting either 150 µl (hydrogels used for immunolabelling and traction force microscopy) or 250 µl (hydrogels used for rheology) of PA solution on plastic cell culture dish lid (#130181, Thermo Fisher Scientific, USA). Smaller cell culture dish lid (#353001, Corning, USA) was placed on top of the PA solution drop for polymerization of 20 min in room temperature. Once polymerized, the PA hydrogels were placed in PBS (#14200-083, Gibco) and stored at +4 °C.

For cell culture, PA hydrogels were cut smaller with puncher (d = 6 mm, #504533, World Precision Instruments, USA) to fit inside the 24-well cell culture insert. To facilitate cell adhesion and growth, the PA hydrogels were coated with Matrigel (#356231, Corning). As proteins cannot spontaneously adhere on PA hydrogels, the hydrogels were first coated with chemical linker 3,4-dihydroxy-L-phenylalanine (L-DOPA) (#D9628, Sigma-Aldrich) according to Wouters et al. (2016). First, 2 mg/ml L-DOPA was dissolved in 10 mM Tris buffer pH 10 (#0497, VWR, USA) for 30 min in dark. The PA hydrogels were incubated with L-DOPA in 24-well plates for 30 min in dark in swing 5 rpm and washed twice with PBS for 10 min. Matrigel was thawed on ice and diluted to 0.1 mg/ml in PBS. This concentration is significantly lower than what was required for Matrigel gel formation (3 mg/ml), resulting in L-DOPA mediated Matrigel protein coating on the PA hydrogel surface. The hydrogels were incubated with the Matrigel solution for 1 h in dark in swing 5 rpm and washed once with PBS. Before cell culture, the hydrogels were sterilized with UV light for 15 min in UV light (UVP CL-1000 Ultraviolet Crosslinker, USA).

### Cell culture

Human embryonic stem cell (hESC) lines Regea08/017, Regea11/013 and Regea08/023 and human induced pluripotent stem cell (hiPSC) line expressing mEGFP tagged beta-actin (actin beta (ACTB) (ID: AICS-0016 cl.184) developed at the Allen Institute for Cell Science (allencell.org/cell-catalog) and available through Coriell (Roberts et al., 2017)) were differentiated and cultured as described previously (Vaajasaari et al., 2011; Viheriälä et al., 2021). Cells were plated on 24-well plate inserts (#83.3932.101, Sarsted, Germany) at a density of 2.5*10^5^ cells/cm^2^. Inserts were coated with 10 µg/cm^2^ Collagen IV from human placenta (#C5533, Sigma) and 1,8 µg/cm^2^ Laminin 521 (LN521, Biolamina, Sweden). RPE cells were maintained at 37 °C in 5 % CO_2_ in DM-media, containing knock-out Dulbecco’s modified Eagle’s medium (#10829018, Gibco, USA) with 15 % knock-out serum replacement (#10828-028, Gibco), 2mM GlutaMax (#35050038, Gibco), 1 % MEM Non-essential amino acids (#11140050, Gibco), 0.1 mM 2-mercapthoethanol (#31350010, Gibco) and 50 U/ml penisillin/streptomycin (#15140122, Gibco). The media was changed every 2–3 days.

PA hydrogels were placed on top of the RPE monolayers after 5 to 7 weeks after plating them on the inserts. First, all the media was removed from the insert. The PA hydrogel was gently placed on top of the cells with tweezers. A spacer was added on top of the PA hydrogel to prevent it from moving or floating. Finally, media was added on top of the hydrogel. During culture, the media was changed every 2–3 days.

For vascular endothelial growth factor (VEGF) analysis, media samples were collected from both basal and apical side of the inserts and stored in -80 °C. Media samples were collected before changing the media (2–3 days after the last media change). VEGF concentration was measured with Human VEGF Quantikine ELISA kit (#DVE00, R&D systems, USA) according to manufacturer’s instructions. To reduce the effect of variation between cell culture inserts, media samples from 2-3 parallel inserts were pooled together to form one sample for ELISA assay, when possible. Media samples were collected altogether from 41 inserts. In the assay each sample was measured 3 times, and an average was calculated for analysis. In this work altogether 2 ELISA assays were made.

Transepithelial electrical resistance (TEER) was measured with epithelial voltohmmeter (EVOM2, World Precision Instruments, USA) with STX2 electrode (World Precision Instruments). TEER was measured from RPE cells cultured with the hydrogel for 2 weeks and from control cells cultured without a hydrogel. The electrode was warmed up in +37 °C cell culture media before and in between the measurements. Each insert was measured 3 times, and an average was calculated for analysis. Empty insert with and without the hydrogel and with cell culture media were measured and subtracted from the averaged TEER values to remove the background resistance. TEER was measured from hiPSC-RPE cells with actin-mEGFP expression by first keeping the cells in RT for 15 min. Each insert was measured 2 times, and the electrode was placed in 1xPBS between each measurement.

### Immunofluorescence staining and confocal imaging

In this work, fixing and immunofluorescent staining of the RPE cells cultured under the hydrogel was done in two different ways. In the first method, the insert membrane was cut with scalpel and the gel was gently removed from the live cells with tweezers. All the following steps were done at room temperature (RT). Cells were fixed in 4 % PFA in PBS 10 min and washed 3 times in PBS for 5 min. Insert membrane was then cut into 2 or 4 pieces with scalpel. Permeabilization was done in 0.1 % Triton-X-100 (v/v) in PBS for 15 min. Blocking was done in 3 % bovine serum albumin (BSA) (w/v) in PBS for 1 h. Primary antibody incubations were done in 3 % BSA (w/v) in PBS for 1 h. Samples were washed 4 times in PBS for 5 min. Secondary antibody incubations were done in 3 % BSA (w/v) in PBS for 1 h in dark. Samples were washed 3 times in PBS for 5 min and mounted between two glass coverslips (18x18 mm, # 474030-9000-000, Zeiss, Germany) in Prolong diamond antifade mountant (#P36961, Molecular Probes, USA). Samples were dried overnight at RT and then stored at +4 °C.

RPE cells were also stained with the hydrogel still attached to cells. First, samples were washed twice with PBS and fixed in 4 % PFA (v/v, #157-8, Electron Microscopy Sciences, USA) in PBS for 30 min at RT. Samples were washed 3 times with PBS for 15 min. Permeabilization was done overnight at +4 °C in 0.5 % Triton-X-100 (v/v, #T8787, Sigma-Aldrich) and 3 % BSA (w/v, #P06-139210, PAN-Biotech, Germany) in PBS. Primary antibody incubation was done in 3 % BSA (w/v) in PBS for 24 h at 4 °C. Samples were washed in 0.5 % Triton-X-100 (v/v) and 3 % BSA (w/v) in PBS three times for 4 h and then overnight at +4 °C. Secondary antibody incubation was done in 3 % BSA (w/v) in PBS for 24 h at +4 °C. Samples were washed in PBS three times for 4 h and then overnight at +4 °C. Samples were stored in PBS at +4 °C. For imaging, the insert membrane and the hydrogel adhered on the cells were cut with scalpel (nro10, Schwann-Morton, UK) and placed on cell culture dish (#130181, Thermo Fisher Scientific). To avoid the sample from floating, insert membrane was adhered on the dish with 3 drops of rapid glue.

Primary antibodies used in this work were: ZO-1 1:50 (#61-7300, Thermo Fisher Scientific), Claudin-19 1:100 (#MAB6970, R&D Systems), Ezrin 1:100 (and 1:200 in Figure 3 – figure supplement 1) ([3C12], #ab4069, Abcam, UK), Ca_V_1.3 1:100 (#ACC-005, Alomone Labs, Israel), Na^+^/K^+^-ATPase 1:200 ([464.6], #ab7671, Abcam), Bestrophin 1:200 (and 1:100 in Figure 3 – figure supplement 1) (#016-Best1-01, Lagen laboratories, USA), CRALBP 1:200 (and 1:500 in Figure 3 – figure supplement 1) ([B2], #ab15051, Abcam), Connexin43 1:200 (#C6219, Sigma-Aldrich), MERTK 1:50 (#H00010461-M01, Abnova, Taiwan), Rac1 1:100 (#610650, BD Transduction Laboratories, USA), PanNa_V_ 1:200 (#ASC-003, Alomone Labs), Vimentin (#ab92547, Abcam). Secondary antibodies were goat anti-rabbit Alexa Fluor 488 1:200 (#A-11008, Invitrogen, USA) and goat anti-mouse Alexa Fluor 568 1:200 (#A-11031, Invitrogen). In addition, goat anti-mouse Alexa Fluor 647 (#A-21235, Invitrogen) was used (Figure 3 – figure supplement 1). Cellular actin was labelled with ATTO 643 Phalloidin 1:100 (#AD643-81, ATTO-TEC, Germany).

Samples with the hydrogel still adhered on the cells were imaged through the hydrogel with Nikon Eclipse FN1 upright spinning disk confocal microscope (Nikon Europe BV, Amsterdam, Netherlands) with Nikon CFI Apo NIR 60x/1.0 water dipping objective (Figure 2A). The light source used in imaging was Spectra X (Lumencor, USA) and the camera iXon 888 EMCCD (Andor, UK). Voxel size was set to x = y = 0.22 µm and z = 0.3 µm and 1024 x 1024 pixel sized images were obtained and saved in .nd2 format. Images were finalized with ImageJ by performing gaussian blur with radius 1.0 and linear brightness and contrast adjustments.

Mounted samples were imaged with laser scanning confocal microscope (LSCM). Zeiss LSM 800 LSCM was used with Zeiss Plan-Apochromat 63x/1.40 oil immersion objective with 488nm, 561nm and 640nm lasers (Figure 1C and Figure 1 – figure supplements 1 and 2 and Figure 3 – figure supplement 1). Voxel size was set to x = y = 0.1 µm and z = 0.15 µm. 1024 x 1024 pixel images were obtained with line averaging 2 and saved in .czi format. Images were finalized with ImageJ by performing gaussian blur with radius 1.0 and linear brightness and contrast adjustments.

### Traction force microscopy

For traction force microscopy (TFM) with the hydrogel cushion (Figure 3), 200 nm fluorescent beads were added to the PA gel (3 % vol/vol, #F8807, Invitrogen) prior to polymerization. RPE cells were cultured under the hydrogel for 2 or 4 weeks. Before imaging, the insert membrane was cut and fixed on cell culture dish (#130181, Thermo Fisher Scientific) with rapid glue with the hydrogel still attached to the live cells. During imaging samples were kept in 5 ml of Ames’ medium (#A1420, Sigma-Aldrich) with 10 mM 4-(2-hydroxyethyl)-1-piperazineethanesulfonic acid buffer (HEPES) (H3375, Sigma-Aldrich), 5 % knock-out serum replacement (v/v, #10828-028, Gibco) and 0.2 % penicillin/streptomycin (v/v, #15140122, Gibco). Cells were kept at +37 °C during imaging. Imaging was done with Nikon Eclipse FN1 upright spinning disk confocal microscope with Nikon CFI Apo LWD 25x/1.1 water dipping objective. Voxel size was set to x = y = 0.52 µm and z = 0.4 µm. 1024 x 1024 pixel timelapse z-stacks were obtained with interval of 5 min or 10 min and duration of 20 min and saved in .nd2 format. For control, after the first 20 min timelapse actin polymerization inhibitor Cytochalasin D (#C2618, Sigma-Aldrich) was added to the imaging media in the concentration of 5 or 10 µg/ml. Imaging was continued for 2-3 more timelapses for 40-60 min, and the last timelapse was used for the image with actin inhibition.

Images of the fluorescent beads were processed in ImageJ by subtracting background (radius 20.0) and adjusting image brightness and contrast linearly. Maximum intensity projections were made from two timepoints with 20 min interval so that only images from near the gel surface were included in the projections. The images were aligned with ImageJ plugin StackReg (rigid body) (Thevenaz et al., 1998). Bead displacement fields were calculated with iterative particle image velocimetry (PIV advanced) in ImageJ (Tseng et al., 2012). 1^st^ pass PIV parameters were set so that the interrogation window size was 64 pixels, search window was set to be interrogation window x 2 (128 pixels) and vector spacing was set to be half from the interrogation window size (32 pixels). For 2^nd^ pass the interrogation window was set to be half from the 1^st^ pass, and for the 3^rd^ pass half from the 2^nd^ pass, so that the final vector spacing was 8 pixels (0.8 µm). In the case of ambiguous vectors in the displacement field, normalized median test (noise 0.20, threshold 5.00) or dynamic mean test (C1 0.20, C2 5.00) were performed. Traction forces were calculated with ImageJ plugin Fourier transform traction cytometry (FTTC) (Tseng et al., 2012). For FTTC, the Poisson ratio of PA gels was set to 0.5, and the gel stiffness was set to 2800 Pa. No regularization was used as our comparison of different λ values did not show a significant effect on the traction magnitudes detected (Appendix 2). For quantification, the output force field text file created by FTTC was used. From each file, an average value was calculated from the force magnitudes (in Pascals). For illustrative purposes, the displayed pixel values of the FTTC stress magnitude maps presented in Figure 3 were adjusted. Original vector plots are shown in Figure 3 – figure supplement 2.

### Phagocytosis assay with the PA hydrogel

The POS particles were extracted as previously described (Vaajasaari et al., 2011). In brief, porcine eyes were obtained from the slaughterhouse, opened and the retinas were removed with tweezers under red light. Retinas were homogenized in 0.73 M sucrose phosphate buffer by shaking and then filtered twice with gauze. POS were separated by ultracentrifugation (Optima ultracentrifuge, Beckman Coulter, Inc., Brea, CA) at 112400 RCF at 4 °C in sucrose gradient. The POS layer was removed, centrifuged at 3000 RCF at 4 °C and stored in 73 mM sucrose phosphate buffer at -80 °C.

POS were thawed and centrifuged at 5400 RCF for 4 min. For labelling, POS particles were mixed with 1 µg/ml ATTO 488 NHS ester (#AD488-31, ATTO-TEC) in 9.5 mM NaHCO_3_ in PBS, incubated for 1 h RT and washed three times with 2800 RCF for 3min centrifugation.

For phagocytosis assay, thin PA gels were constructed on glass coverslips with POS particles incorporated on the hydrogel surface. First, both the top (d = 13 mm, VWR, USA) and bottom coverslips (22 x 22 mm, No. 1.5H, Marienfeld, Germany) for PA gel construction were washed in 2 % HellmaneX III (Hellma, Germany) solution in MQH_2_O for 30 min in a sonicator, rinsed with excess water and ethanol and dried with compressed nitrogen. The bottom coverslips were activated for the PA gel to adhere to it. Bind-silane (3-(Trimethoxysilyl)propyl methacrylate, #M6514, Sigma-Aldrich) and glacial acetic acid were diluted with 95 % ethanol in final concentrations of 0.3 % v/v and 5 % v/v, respectively. The solution was let to react for 3 min on the bottom coverslips. Next, the bottom coverslips were washed with ethanol and dried with compressed nitrogen. The top coverslips were incubated with the ATTO labelled POS particles for 1 h at RT and washed by dipping the coverslip into a beaker with PBS. Acrylic acid N-hydroxysuccinimide ester (NHS-AA, #A8060, Sigma-Aldrich) was diluted in water-free dimethyl sulfoxide (DMSO, #D2650, Sigma-Aldrich) in 50 mg/ml and further diluted to 0.1 mg/ml in 9.5 mM NaHCO3 in PBS. NHS-AA solution was let to react on the top coverslips for 30 min at RT, and after the incubation washed by dipping the coverslip into a beaker with PBS. During the NHS-AA incubation, polyacrylamide solution for E = 2.8 kPa hydrogel was mixed and degassed as described above. 200 nm fluorescent beads were added to the PA gel (2 % v/v, F8810, Invitrogen) prior to polymerization with TEMED and APS (10 % w/v). 13 µl of PA solution was pipetted on bind-silane activated bottom coverslip, and d = 13 mm top coverslip with the POS and NHS-AA coating was added on top. Gels were let to polymerize 15 min in dark in RT. After polymerization gels were stored in PBS at +4 °C overnight. The next day coverslips were gently removed with scalpel. For imaging, PA gels on coverslips were placed on live cell imaging chambers (#SC15022, Aireka Cells, China). Control gels were constructed similarly as described above. For controls with NHS-AA on the surface, NHS-AA was incubated on clean top coverslip at 0.1 mg/ml in 9.5 mM NaHCO3 in PBS for 30 min and rinsed with PBS.

For live cell imaging, cellular actin was labelled with SiR-Actin kit (#CY-SC001, Cytoskeleton Inc., USA). hESC-RPE cells (cultured for 10 to 18 weeks after plating on inserts) were incubated at +37 °C in 5 % CO_2_ in DM-media containing 25 pmol Sir-Actin for 3 h. Cells were washed once with DM-media. Insert membrane was cut with scalpel and placed on the PA gel in a live cell imaging chamber (#SC15022, Aireka Cells, China) with cells facing down to the POS. A slice anchor (Warner Instruments) and a spacer were added on top of the insert to prevent it from floating. Imaging was done in Hibernate-A media (#A1247501, Gibco) with 10 % fetal bovine serum (FBS, # 10500064, Gibco). For controls without serum, just Hibernate media was used. Zeiss LSM 780 LSCM was used with Zeiss LD C-Apochromat 40x/1.10 water immersion objective with 488 nm, 561 nm and 640 nm lasers. Voxel size was set to x = y = 0.1 µm and z = 0.37 µm. 1024 x 1024 pixel timelapse z-stacks with interval of 10 min and duration of 20 min were obtained and saved in .czi format. Cells were kept at +37 °C during imaging. From each insert, 3 separate positions were chosen and imaged alternately. For control, 10 µg/ml Cytochalasin D (#C2618, Sigma-Aldrich) was added to the medium during imaging without touching the sample.

Cellular tractions were analyzed from the bead images as was described in section 5.4. Images of POS and cellular actin were finalized with ImageJ by performing gaussian blur with radius 1.0 and linear brightness and contrast adjustments. To quantify the traction stress magnitudes around POS particles, 3 circular regions with diameter of 100 pixels (10.4 µm) were chosen from each traction stress magnitude image, which illustrate the change in traction stress during cell actin cytoskeleton relaxation with Cytochalasin D. For control, the same areas chosen for the 1^st^ image were used in the 2^nd^ image, thus creating randomized control areas within the same dataset. The intensity of the areas chosen was then measured from the 32-bit FTTC traction stress magnitude images, where the pixel values represent the traction stress magnitude in Pascals. For illustrative purposes, the displayed pixel values of the stress magnitude maps in Figure 4C and Figure 4E have been adjusted. Original vector plot of the stress magnitude map in Figure 4E is shown in Figure 4 – figure supplement 1.

### Rheology

A rheometric tack adhesion test was performed to quantify the RPE-hydrogel adhesion using Discovery Series Hybrid Rheometer II from TA Instruments. First, the insert membrane and the hydrogel adhered to the live cells were cut from the insert with a scalpel. The insert membrane was fixed on a 22 x 22 mm glass coverslip (#0107052, Marienfeldt, Germany) with rapid glue, and the coverslip was taped on the bottom parallel plate of the rheometer. Then, the hydrogel was glued on the probe (top 8 mm parallel plate). A holding period of 2 min with compressive force of 0.1 N was applied. Then, the hydrogel was separated from the RPE cells by moving the probe upwards at a constant linear rate of 10 µm/s. The measurement was set to last for 360 s. As a result, a force-distance curve was drawn in Origin (2019b). At the end of the measurement, the force remained constant as the probe was moving upwards with the detached hydrogel still adhered to it (baseline, illustrated s a red line in Figure 2B). Adhesion force was defined from each curve by subtracting the baseline force from the lowest peak of the curve (detachment point). Adhesion energy was measured graphically from each curve by calculating the area between the curve and the baseline force. As a control, cells were treated with 10 µg/ml Cytochalasin D (#C2618, Sigma-Aldrich) for 1 h at +37 °C in 5 % CO_2_. For a second control, cells were first treated with Cytochalasin D and then fixed with 4 % PFA (v/v, #157-8, Electron Microscopy Sciences, USA) in PBS for 30 min at RT.

### Statistical testing

Statistical tests were performed using the IBM SPSS statistics for Windows, version 28 (IBM Corp.) with a significance level of P < 0.05. Normality was tested using Kolmogorov-Smirnov and the Shapiro-Wilk tests. Graphs were made with Origin (2019b). In the graphs, median is illustrated with a horizontal line and mean with a white box. Boxes consist of 25-75% of the sample range, and the line is range within 1.5IQR. Schematics and final figures were assembled in Adobe Illustrator.

## Acknowledgements

We thank Outi Heikkilä, Outi Melin and Hanna Pekkarinen for their contributions with the cell culture work. We thank Jenni Keränen for technical assistance. We thank Jens Möller and Nonappa Nonappa for the discussions regarding this work. The authors acknowledge the Biocenter Finland (BF) and Tampere Imaging Facility (TIF) for the service.

## Abbreviations

AA: Acrylamide
APS: Ammonium persulfate
BIS: N,N’-Methylenebisacrylamide
BSA: Bovine serum albumin
ECM: Extracellular matrix
mEGFP: Monomeric enhanced green fluorescent protein
FTTC: Fourier transform traction cytometry
HEPES: 4-(2-hydroxyethyl)-1-piperazineethanesulfonic acid
hESC: Human embryonic stem cell
hiPSC: Human induced pluripotent stem cell
IPM: Interphotoreceptor matrix
L-DOPA: 3,4-dihydroxy-L-phenylalanine
PA: Polyacrylamide
PBS: Phosphate buffered saline
PFA: Paraformaldehyde
PIV: Particle image velocimetry
POS: Photoreceptor outer segment
RPE: Retinal pigment epithelium
TEER: Transepithelial electrical resistance
TFM: Traction force microscopy
VEGF: Vascular endothelial growth factor

