## Supplemental materials for "Biomimetic hydrogel platform reveals active force transduction from retinal pigment epithelium to photoreceptors"

### Figure supplements

**Figure 1 – figure supplement 1. Immunofluorescence staining of RPE markers.**

Representative confocal images of RPE markers in 7-week-old hESC-RPE cells cultured with PA hydrogels for 2 weeks (controls cultured without a hydrogel), labeled with immunofluorescence staining after removing the hydrogel. XY sections: maximum intensity projections, YZ sections: maximum intensity projections of 5 images with location indicated as white line in the XY section. Scalebars 10 µm.

**Figure 1 – figure supplement 2. Topography of the hydrogel surface might affect the RPE.**

(A) Maximum intensity Z-projections of Ezrin and Phalloidin (actin) of hESC-RPE cells cultured with the conventional method (top row) and with the PA hydrogel polymerized between two plastic surfaces (middle row) or between two glass surfaces (bottom row). Differences in cellular actin organization indicate that the topography of the hydrogel surface may affect the cells. (B) Schematic illustrating the topography construction on PA hydrogel surface. SU-8 master was constructed with contact photolithography with roughly photoreceptor-sized pillars (d = 3–5 µm, h = 5 µm). SU-8 was then used to mold a PDMS structure, which was then used to cast the topography on a thin PDMS layer on glass coverslip. This PDMS coverslip was then used to cast the topography on the PA hydrogel surface. (C) Confocal image of a PA hydrogel labelled with fluorescein-o-acrylate with the photoreceptor-mimicking topography. Scalebars 10 µm. (D) Schematic illustration of RPE cells cultured under the PA hydrogel with photoreceptor-mimicking topography. Ideally RPE's apical side microvilli could locate inside the wells on the gel surface to resemble the tight, interlocked structure of RPE-photoreceptor interaction seen *in vivo*.

**Figure 3 – figure supplement 1. Characterization of the actin-mEGFP hiPSC-RPE cells.**

(A) Representative confocal images of RPE markers in 10–11-week-old hiPSC-RPE cells with actin-mEGFP expression, labelled with immunofluorescence staining. Maximum intensity projections, scalebars 20 µm. (B) Transepithelial electrical resistance (TEER) measured from 11 and 13 -week-old hiPSC-RPE cells with actin-EGFP expression.

**Figure 3 – figure supplement 2. Vector plots of the FTTC data.**

Tractions shown as vector plots of the FTTC data presented in Figure 3D as stress magnitude maps. Force magnitude is illustrated with color in Pascals. Scalebars 50 µm.

**Figure 4 – figure supplement 1. Vector plots of the FTTC data.**

Vector plot of the FTTC data from Figure 4E stress magnitude maps. Left: Composite image of the POS particles (green, maximum intensity Z-projection) and vector plot of the tractions, right: the vector plot. White circles illustrate the areas (d = 10 µm) around the POS particles used in analysis. Force magnitude is illustrated with color in Pascals. Scalebars 10 µm.

**Figure 4 – figure supplement 2. Traction stresses without POS particles.**

As a control, we used PA gels without POS particles, and the traction forces were on a scale of tens of pN/$\mu m^2$. We constructed hydrogels with and without NHS-acrylic acid supplementation (+NHS-AA and -NHS-AA). RPE cells were supplemented with serum during imaging (+ 10 % FBS, -NHS-AA n = 18 images, 3 inserts and +NHS-AA n = 18 images, 2 inserts) or were imaged without serum (- FBS, n = 13 images, 3 inserts).

### Figure 1 – figure supplement 1

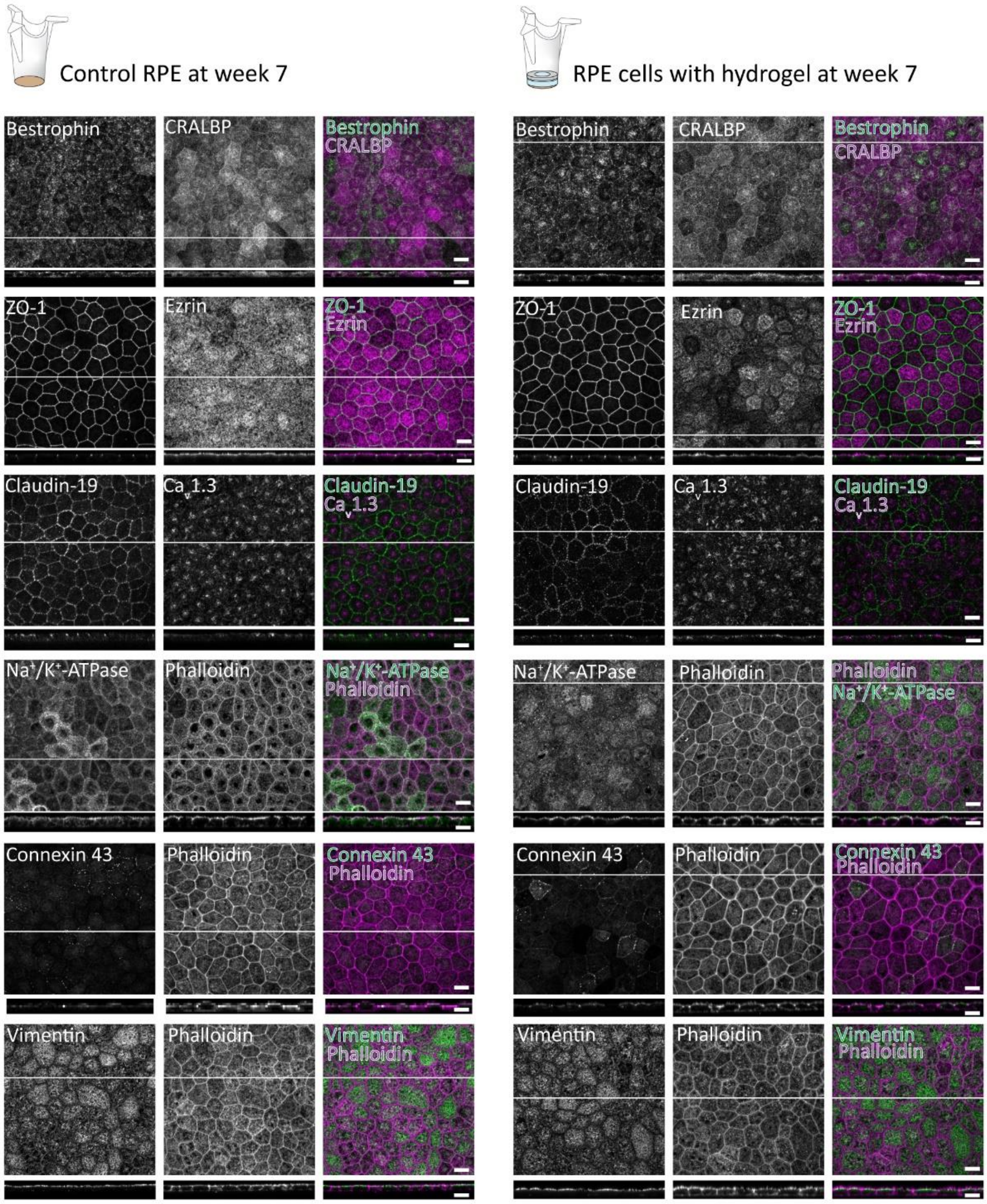

### Figure 1 – figure supplement 2

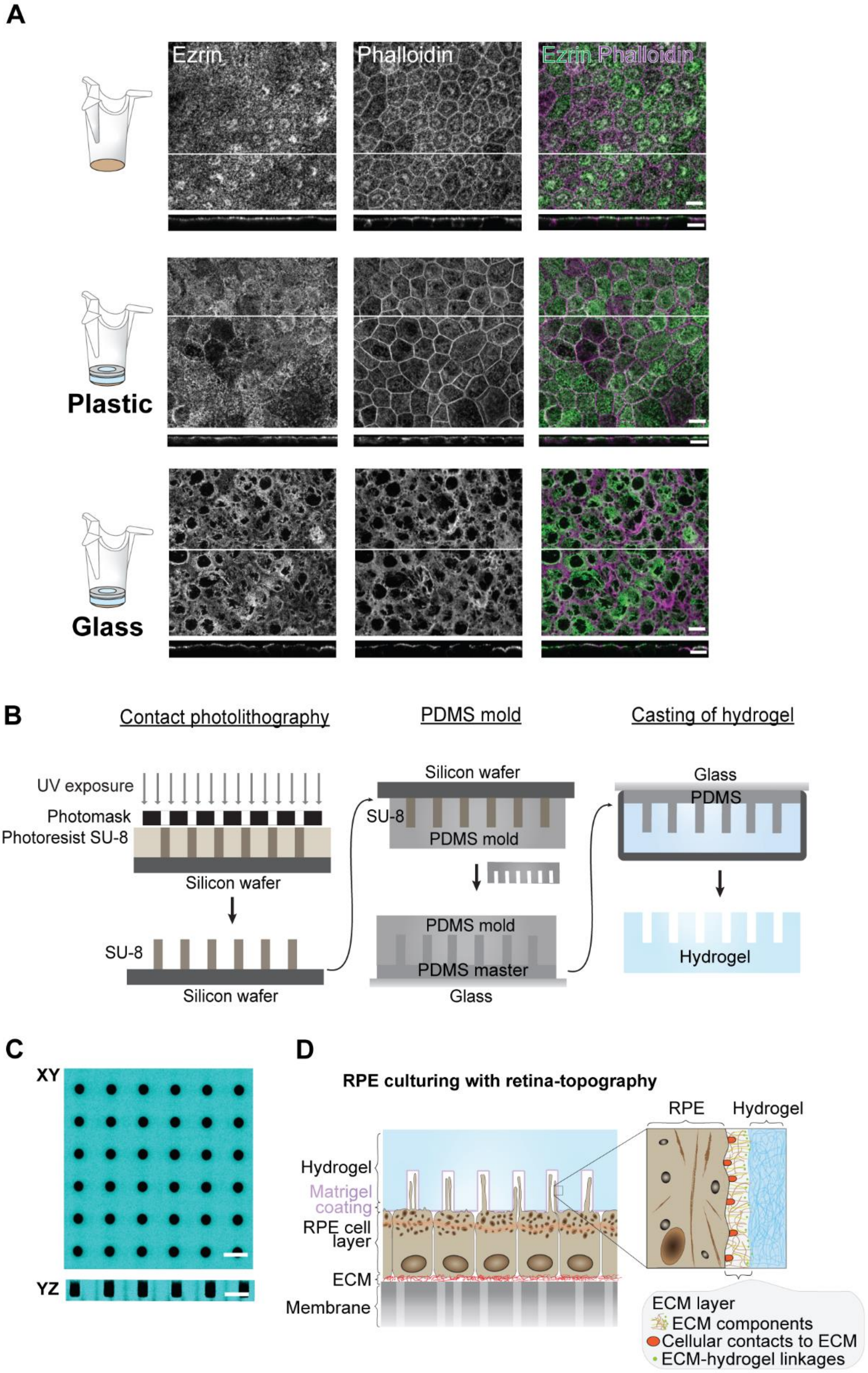

### Figure 3 – figure supplement 1

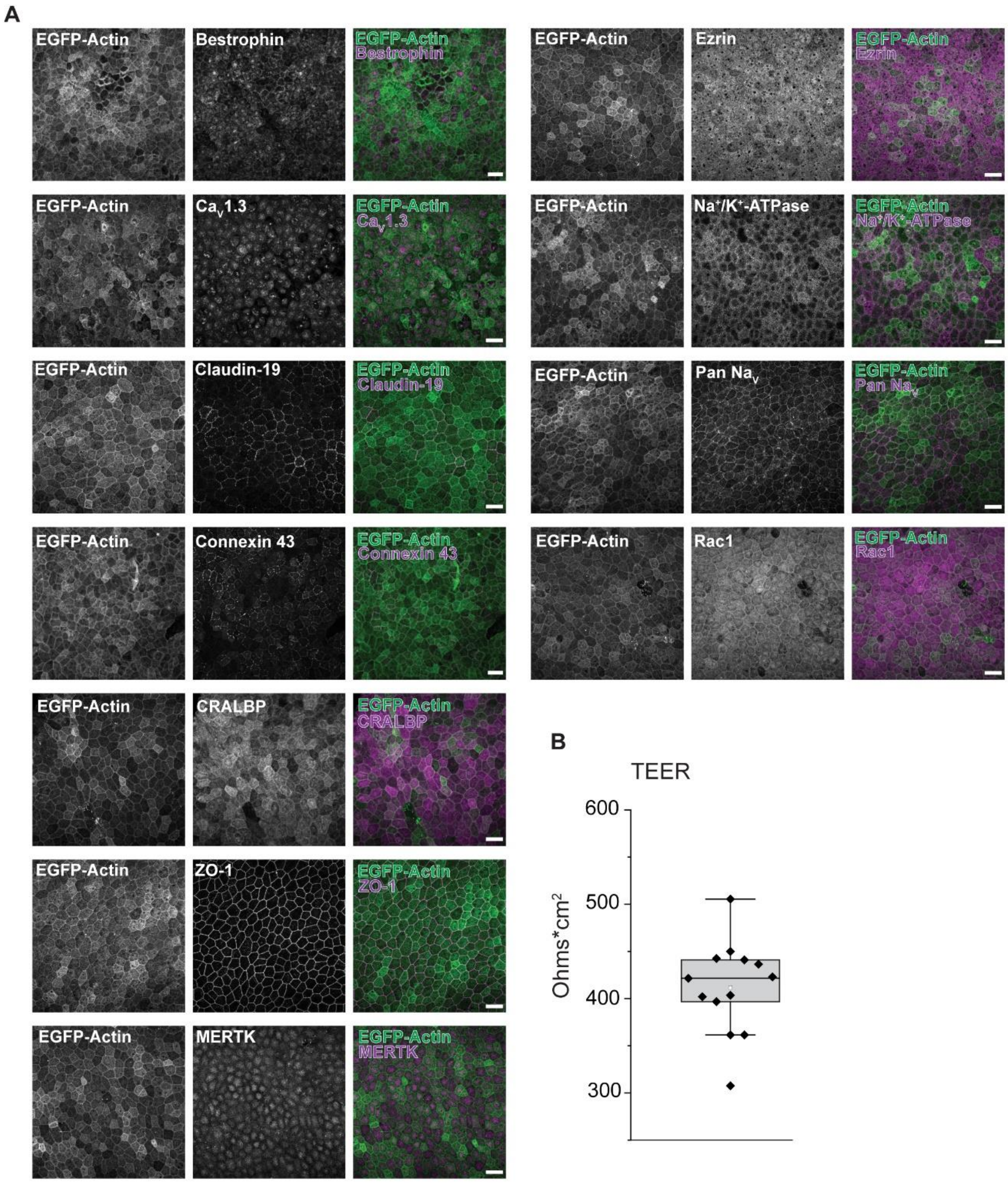

### Figure 3 – figure supplement 2

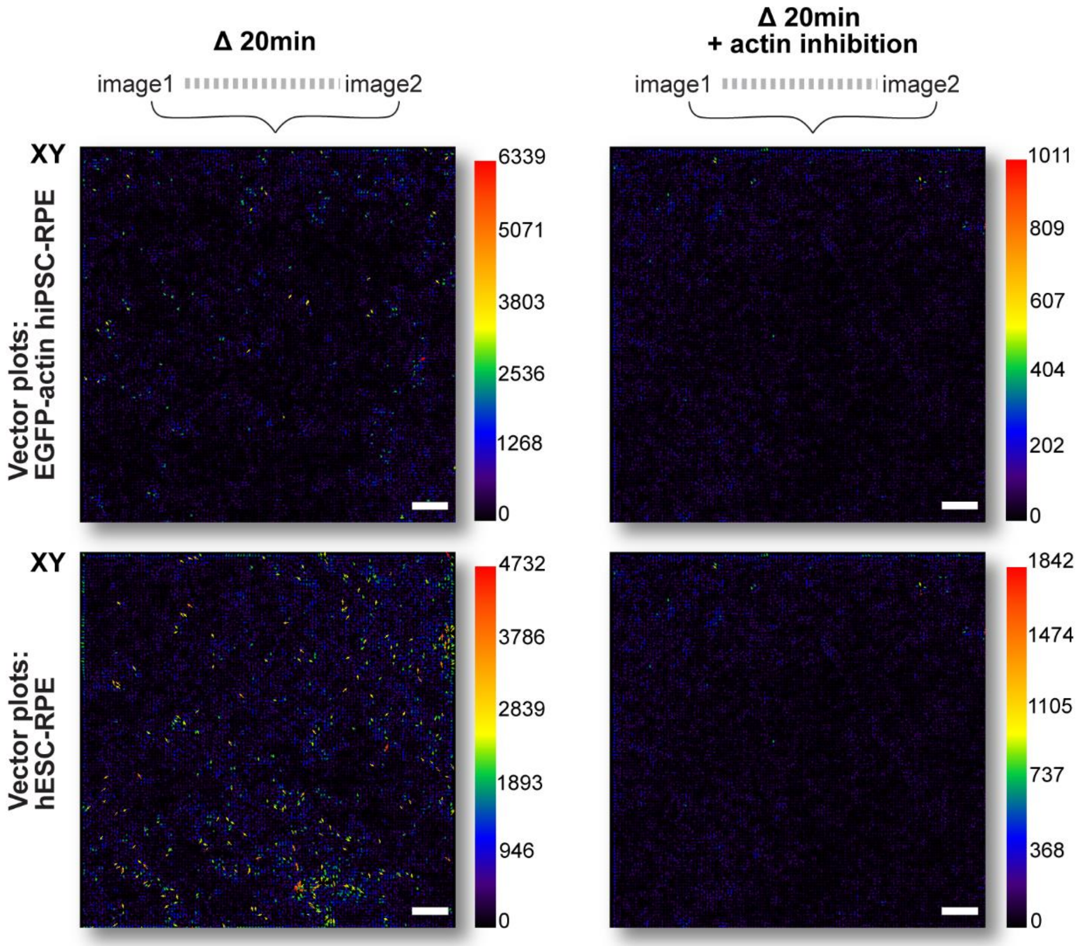

### Figure 4 – figure supplement 1

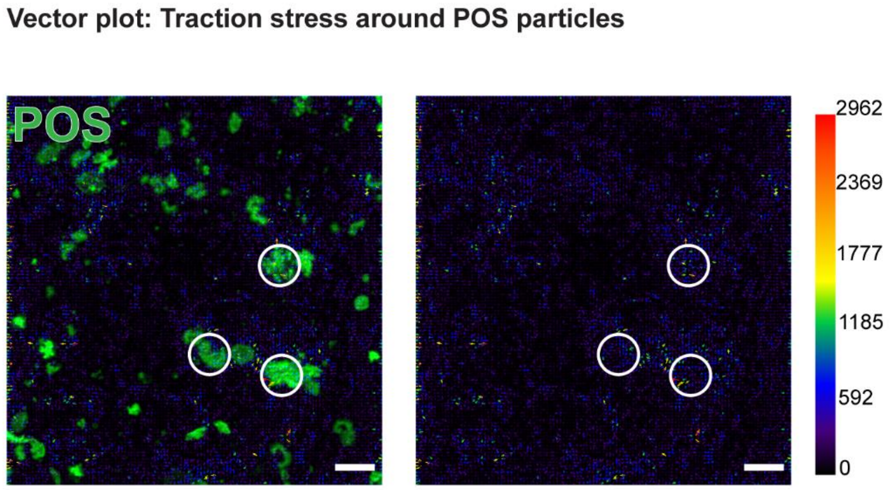

#### Figure 4 – figure supplement 2

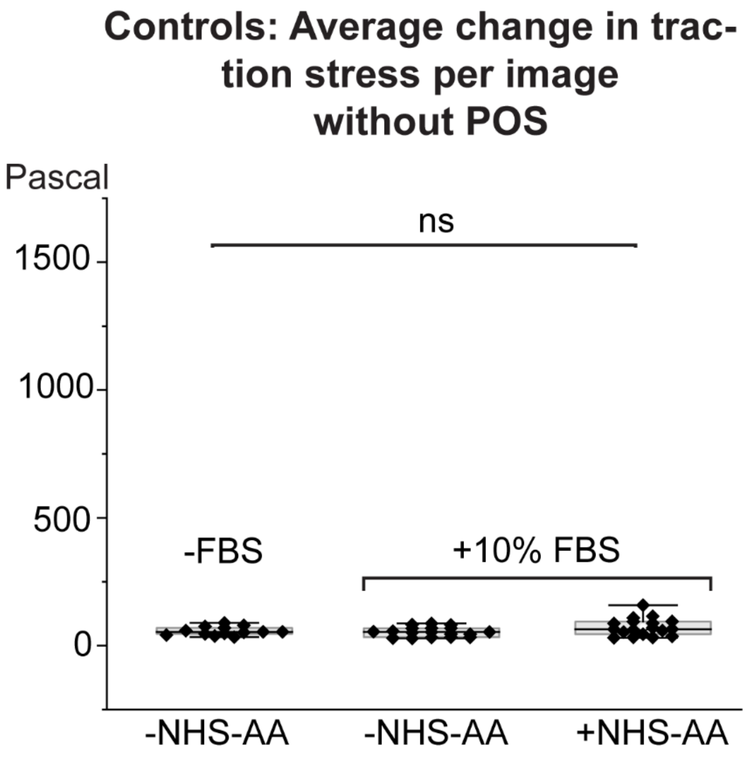

### Appendix 1. Topography construction on PA gel surface.

#### Photomask fabrication and photolithography

Photomask design was done using AutoCAD® (Autodesk, 2019). Roughly photoreceptor-sized pillars (d = 3 – 5 µm) were drawn in 5 mm x 5 mm area. The photomask was fabricated in a cleanroom facility (Class ISO 6, UV-protected). Fabrication of a cs hardmask blanc (Clean Surface Technology CO., Japan, CBL5009Du-AZ1500) was done using µPG501 Conversion Job Manager (Version 1.8.3, Heidelberg instruments). First, the AutoCAD .dxf file was converted to HIMT-mode and exposure of a mask blanc was done with exposure time 22 ms, DEFOC = -2. After that, the mask blanc was unloaded from µPG501 and developed in 1:4 mixture of AZ351B/dH2O (AZ Electronic Materials, Germany, 10054724960) for 1 min, washed with dH2O and dried with pressurized nitrogen. Mask was etched in CHROME ETCH 18 (OSC OrganoSpezialChemie Gmbh, 79600316) for 1 min, washed with dH2O and dried with pressurized nitrogen. Finally, the mask was exposed in UV-F 400 Vitralit light box (Panacol-Elosol Gmbh) to UV light (intensity 25 mW/cm2) for 1 min and developed again in the previously made AZ351B/dH2O solution. Mask was washed with dH2O and dried with pressurized nitrogen.

For SU-8 master, a silicon wafer (d = 100 mm, ID#452, University wafer) was treated with O2 plasma for 2 min in reactive ion etching RIE (Advanced vacuum vision 320 RIE). After that, the wafer was coated with 5 µm layer of SU-8 5 photoresist (Y131252 0500L 1GL, Mi-crochem) in spin coater (Laurell, model WS-650HZB-23NPPB). Spin-coating was done in three steps: First 15 s in 500 rpm (acceleration 100), then 40 s in 3000 rpm (acceleration 250), and lastly 10 s in 100 rpm (acceleration 500). After the spin-coating, the wafer was baked on top of a copper plate that was placed on a hotplate. Temperatures were ramped so that temperatures of the copper plate were measured with a thermometer (Tenma, 72-7715). The wafer was baked first on a 65 °C copper plate for 1 min, and then on a 95 °C copper plate for 3 min (Torrey pines scientific Inc., models HS60 and HS61). After that, the wafer and the copper plate were removed from the hotplate. The wafer was cooled down on the copper plate until temperature was below 40 °C.

Exposure of the wafer was done with OAI mask aligner (Optical Associates Inc., model J500/VIS, intensity 12 MW/cm2 at 436 nm wavelength, i-line filter installed). Wafer was placed in contact with the photomask and exposed for 20 s. After exposure, the wafer was heated again in 65 °C for 1 min and 95 °C for 2 min with temperature ramping as described above. Once the wafer had cooled down below 40 °C it was developed in MR-DEV600 developer (Microchem) for 1 min, rinsed with isopropanol and dH2O and dried with pressurized air. If the wafer was not properly developed, the developing was repeated. SU-8 5 patterns on the silicon wafer surface were observed with microscope. Stylus profiler (DektakXT, Bruker) was used to measure the height of the SU-8 5 photoresist layer. After removing the SU-8 master from the SI-cleanroom a final hardbake was done at 150 °C for 15 min. SU-8 master was slowly cooled down in the oven overnight to avoid SU-8 layer from cracking.

#### PDMS molding

The SU-8 master structure was used to cast poly(dimethylsiloxane) (PDMS) structures. First, SU-8 master was treated with Trichloro(1H,1H,2H,2H-perfluorooctyl)silane (PFOCTS, Sigma, 448931-106). PFOCTS treatment was done in chemical hood by keeping SU-8 master and 20 µl of fluorosilane on a glass coverslip in plastic container overnight. As PFOCTS evaporates, it creates superhydrophobic and anti-fouling surface on SU-8 master, which helps removal of PDMS after molding.

For PDMS molding, silicon elastomer was weighted (Sartorius AY612) and mixed with curing gent of 10 % of the base weight of elastomer (Dow, Sylgard 184 silicon elastomer kit, 04019862). Solution was poured on top of the SU-8 master on a petri dish and de-gassed with vacuum until it was clear of bubbles. PDMS was heated in oven (Binder, model FD 23) at +60 °C for 6 h. PDMS master was then peeled off from SU-8 master with tweezers.

These PDMS master structures were then used to mold thin PDMS layers with the topography on glass coverslips. First, PDMS masters were treated with PFOCTS similarly as with SU-8 masters. Before PFOCTS coating PDMS masters were treated with O2 plasma in 0.3 mbar with 50 W power for 27 s (Diener Pico Plasmacleaner). PDMS solution was made again by mixing silicon elastomer and curing agent and degassing the solution, as described above. PDMS-PDMS casting was done on a glass coverslip so that a drop of degassed PDMS solution was placed on a 22 x 22 mm glass coverslip (Marienfeld superior, 0107052), and PDMS mold was stamped on top of that droplet. Coverslips were heated in oven for 6 h. After heating the PDMS stamp was removed from the coverslip. As a result, the coverslip had only a thin layer of PDMS on top, but still contained the topography structure. These PDMS-coverslips were then treated with O2 plasma 100 W in 0.3 mbar for 5 min (Diener Pico Plasmacleaner) to make the PDMS harder and more durable.

#### PA hydrogel casting

PA hydrogels with patterned topography on the surface were fabricated similarly as described previously. However, polymerization of the hydrogels was done between a 5 cm petri dish (Thermo Fisher Scientific, 130181) and the PDMS-coverslip, which were separated with h = 0.5 mm metallic spacer. For confocal imaging, fluorescein-o-acrylate (Sigma-Aldrich) was added to the PA solution in a concentration of 0.25 mg/ml. Gels were let to polymerize 15 min in room temperature until flooded with 1xPBS and stored in +4 °C. Imaging was done with Nikon A1R+ laser scanning confocal microscope with Nikon CFI Plan Apo IR SR 60x/1.27 water immersion objective with 488 nm laser. Voxel size was set to x = y = z = 0.1 µm and 1024 x 1024 pixel images were acquired and saved in .nd2 format.

### Appendix 2. Comparison of FTTC regularization factors.

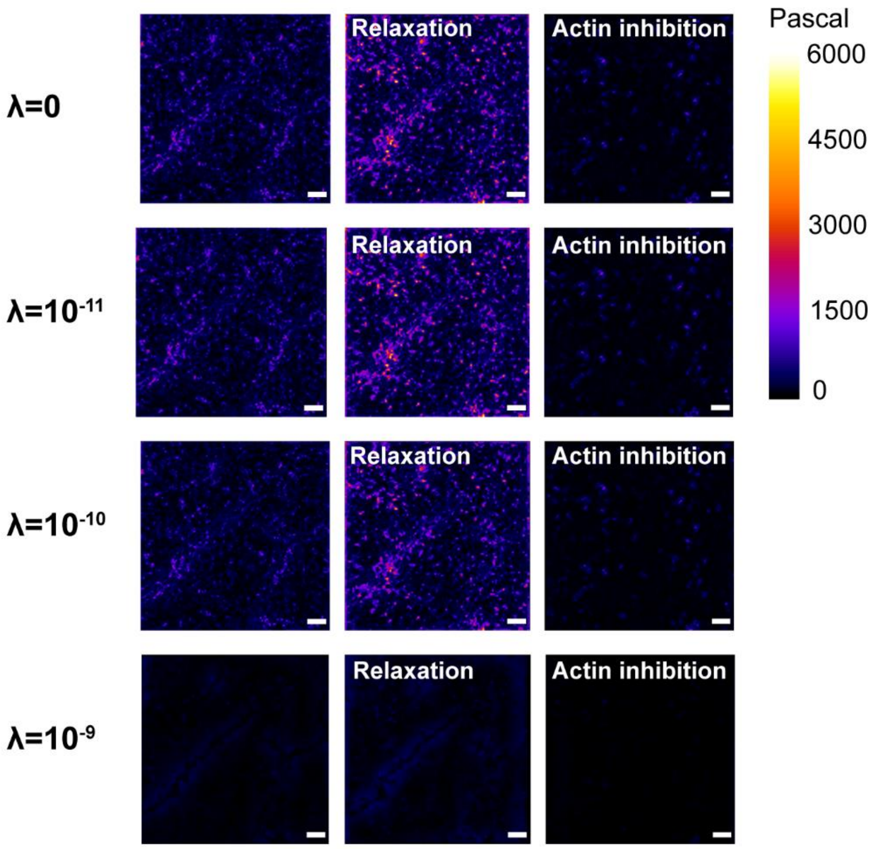


FTTC is sensitive to noise in the bead displacement field. We compared our data without regularization ($\lambda = 0$) and with regularization ($\lambda$: $10^{-11}$, $10^{-10}$, $10^{-9}$) in FTTC plugin in ImageJ. Scalebars 10 µm.
